# Prostaglandin E2 facilitates reciprocal crosstalk between intestinal smooth muscle tissue and epithelium in tumor development

**DOI:** 10.64898/2026.06.12.731555

**Authors:** Rouan Yao, Lilith Sian Lee, Marion Duchemin, Alberto Díez-Sánchez, Andrew B. Single, Anne Marstad, Unni Nonstad, Animesh Sharma, Håvard Takle Lindholm, Lars Hagen, Mesut Bilgen, Menno J. Oudhoff, Mara Martín-Alonso

**Affiliations:** Department of Clinical and Molecular Medicine (IKOM), Norwegian University of Science and Technology (NTNU), Trondheim, Norway; Cellular and Molecular Imaging Core Facility (CMIC), Norwegian University of Science and Technology (NTNU), Trondheim, Norway; Central Administration, St. Olavs Hospital, Trondheim University Hospital, 7030, Trondheim, Norway; Proteomics and Modomics Experimental Core Facility (PROMEC), Norwegian University of Science and Technology the Central Norway Regional Health Authority, Trondheim, Norway; Department of Pathology, Oslo University Hospital-Rikshospitalet, Oslo, Norway; Lipidomics Core Facility, Danish Cancer Institute, Copenhagen, Denmark; Department of Health Sciences, Carleton University, Ottawa, Ontario, Canada

## Abstract

Colorectal cancer arises from intestinal epithelial cells and is driven by the accumulation of aberrant signaling events that disrupt intestinal stem cell (ISC) homeostasis. While stromal regulation of the ISC niche is increasingly recognized, the contribution of the underlying/adjacent smooth muscle tissue (SMT) remains poorly characterized. In particular, the potential crosstalk between normal or tumoral epithelium and the SMT is largely unexplored.

Here, we investigate the ability of the SMT in modulating normal and tumorigenic epithelial biology, and the impact of the tumoral epithelium on the underlying SMT. Mechanistically, by using imaging analyses, transcriptomics, and mass spectrometry, we identify the SMT-derived factor Prostaglandin E2 (PGE2) as the key driver of epithelial dedifferentiation, promoting YAP transcriptional activation, organoid enlargement, and spheroid morphology in intestinal epithelial organoids following SMT supernatant exposure. Conversely, we show that non-invasive tumoral epithelium induces significant changes in the underlying SMT. Tumor-associated SMT shows structural remodeling, inflammation, and reduced muscle fitness. Furthermore, we detected increased prostaglandin signaling, including upregulation of PGE2 synthase *Ptgs1* (COX1), in the SMT underneath the epithelial tumor. Our results reveal a communication between the tumoral epithelium and the underlying SMT prior to metastasis, which could be a facilitating step for tumoral extramural progression. Taken together, these findings highlight a novel tumor-stroma interaction.

## Introduction

Much of the effort in understanding intestinal tumor development and evolution has focused on tumor cells arising from the accumulation of mutations in epithelial cells. Such studies have provided key insights into aberrant signaling pathways in the intestinal epithelium, including the Wnt/β-catenin, MAPK/ERK, BMP/SMAD4, and p53 pathways, which drive more aggressive proliferation of intestinal stem cells (ISCs) and lead to cancer[1].

Epithelial homeostasis, understood as the balance between ISCs proliferation and differentiation, is carefully regulated by niche growth factor gradients, importantly, WNT family members, which maintain ISC stemness, and members of the bone morphogenic protein (BMP) family and their antagonists, which drive maturation[2]. These niche factors control ISC number, proliferation, and injury responses through their complex spatial distribution and molecular expression profiles[3]. Furthermore, alterations in the signaling pathways responding to these niche factors, such as the classic, constantly activated WNT signaling in epithelial cells, *via* mutations in APC, are necessary initial steps in epithelial transformation that will lead to tumor formation[1]. ISC niche factors are provided by other epithelial cells and, as recently shown, by adjacent stromal cells[3–6].

Due to the importance of these niche factors in ISC regulation and cancer development, many efforts have been made to categorize the different populations of peri-epithelial stromal cells and their niche-supporting capabilities. Recent studies have identified multiple fibroblast types as part of these stromal populations, classified based on their distance to the epithelium and growth factor expression profile[3, 4, 7]. Specifically, recent investigations have revealed that specific PDGFR-a (Platelet-derived growth factor receptor-alpha)+ cell subtypes underlying the epithelium can contribute to epithelial tumorigenesis by paracrine signaling[7]. One of the paracrine effectors secreted by these PDGFRa+ cells is Prostaglandin E2 (PGE2), a small lipid mediator with pleiotropic effects over many organs and tissue types due to a range of PGE2 receptors, EP1-4, that have differential ligand affinities and stimulate a variety of downstream effector cascades[7–9]. In the normal intestinal epithelium, PGE2 is shown to play a protective role by stimulating mucin production by goblet cells, as well as inducing the differentiation of wound-associated epithelial (WAE) cells, which help to maintain epithelial barrier integrity upon injury[10–12]. However, PGE2 can also act as an active paracrine driver of early tumorigenesis, through EP4-dependent YAP activation and expansion of Sca1+ reparative stem-like cells[7, 13], and aid in further tumor support and progression by inducing tumor cell growth, survival and expansion, angiogenesis, and immune evasion through multiple signaling pathways[14, 15]. PGE2 downstream signaling includes PI3K/Akt and MAPK/ERK pathways, β-catenin stabilization, and HIF1α-associated responses, among others[16].

Underlying the intestinal epithelium, there are three concentric layers/rings of involuntary contracting smooth muscle. Until recently, the role of this intestinal smooth muscle tissue (SMT) was thought to be limited to the mediation of peristaltic and segmented intestinal movements, which serve to mix digestive enzymes with food, as well as shuttle food along the intestines. Importantly, PGE2 also plays a role in the modulation of muscle contractility, proliferation, and differentiation, suggesting that it is produced in abundance in all muscle tissues, including intestinal smooth muscle[17–21]. In recent years, our work[22] and that of others[23, 24] have shown that the SMT plays a previously unsuspected role in the ISC niche, in the supply of fundamental niche growth factors, in particular BMP antagonists, thereby influencing epithelial stemness/differentiation in homeostasis. Specifically, we have shown that SMT modifications, such as the loss of expression of the matrix metalloproteinase MMP17, impair mucosal healing and decrease the epithelial ISCs pool[22].

Furthermore, the SMT constitutes a natural barrier to epithelial tumor metastasis, where tumor dissemination/progression commonly involves invasion through the SMT[25, 26]. Despite this, epithelial-SMT crosstalk both in homeostasis and throughout epithelial transformation, tumor growth, and metastasis remains profoundly understudied.

Here, we identify the SMT as a source of critical factors in epithelial tumorigenesis and characterize the bidirectional communication between the epithelium and the SMT, with a particular focus on how this interaction is modulated in the presence of an epithelial tumor. By exposing intestinal organoids to SMT-derived supernatants, we found that SMT secretes factors that increase organoid size and reduce epithelial differentiation. We identified the SMT-derived molecule PGE2 as a key driver of this phenotype, acting through the PGE2-EP4-YAP/HIF1α axis. Furthermore, our tissue analyses show that the SMT is profoundly modified by non-invasive adenomas, showing increased PGE2-synthase COX1 expression, inflammation, and downregulation of muscle contractile markers.

In sum, we demonstrate that (i) the SMT is a major source of PGE2, a tumor-initiating, promoting, and supporting molecule, and (ii) the intestinal SMT is extensively remodeled underneath the epithelial tumor. Together, these findings suggest that tumor-associated SMT undergoes changes that confer a selective advantage to tumor growth and progression, potentially priming it for transmural progression in later tumoral stages.

## Results

### Muscle supernatant (MSN) exposure induces morphological changes in WT and tumor-derived organoids, resulting in heightened YAP-Sca1 activation in WT organoids

Previous work from our group showed that exposure of organoids to intestinal smooth muscle-conditioned medium (muscle supernatant, MSN) for four days renders wild-type (WT) healthy organoids more proliferative, less differentiated, morphologically resembling tumor organoids (i.e. large spheroids instead of budding organoids), and with YAP transcriptional regulator activated[22]. YAP activation has been previously shown to play a role in tumor initiation driven by *Apc* deficiency[27, 28]. Furthermore, we demonstrated that the smooth muscle tissue (SMT) is a source of BMP antagonists, which will limit levels of SMAD4 in the epithelium, a key tumor suppressor gene usually downregulated in colorectal cancer (CRC). Collectively, as these observations suggested a possible implication of the SMT in epithelial tumor regulation, we sought to determine the nature of MSN-epithelial interaction in WT and tumor organoids.

Apc^Min/+^ mice are widely used for their autochthonous tumor-initiating capacity. Three to four-month-old Apc^Min/+^ spontaneously developed tumors in the intestine that ranged from small polyps to well-formed adenomas. Here, we exposed WT and Apc^Min/+^ tumor-derived organoids to MSN and analyzed early responses. After MSN exposure, organoids increased their size when compared to untreated control organoids, becoming significantly bigger 48 hours after exposure (Figure 1a,b and Figure S1a). For the following days, WT organoids grew significantly larger upon MSN exposure and shifted their morphology, showing a higher percentage of spheroids (Figure S1a,b). Tumor organoids maintained their spheroid-like morphology as expected, but grew significantly larger in response to MSN (Figure 1a,b, and Figure S1a). This difference in size disappeared at day 3 (Figure S1a, c). Shape and size changes were analyzed using automated image analyses (see methods).

**Figure 1.**
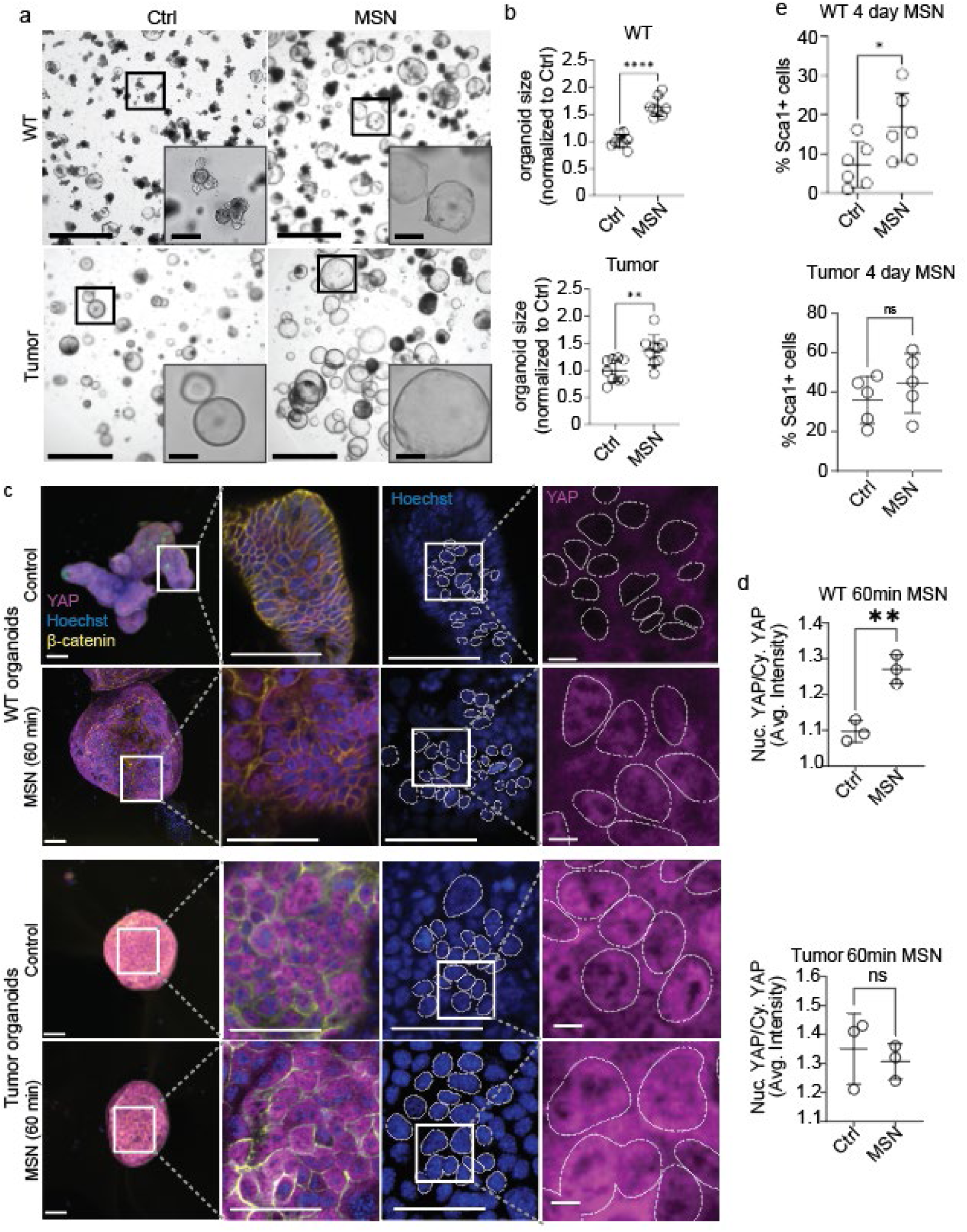
Smooth muscle-derived factors induce a morphological shift in both wild-type and tumor organoids and YAP activation in wild-type intestinal organoids. A. Representative bright-field images of wild-type (WT) and Apc^Min/+^-derived tumor organoids 48h after treatment with media supplemented with smooth muscle-derived supernatant (MSN) or regular ENR or EN for tumoroids (here referred to as Control media). Scale bars = 1250µm and 200µm in zoomed insets. B. Graphs show organoid size in culture by using automated image analysis. Points represent mean values from each biological replicate (total of n=8 for WT organoids and n=9 for tumor organoids, 3-4 wells quantified per replicate), each treated with Control or MSN media. Data were analyzed with paired t-tests. Relative organoid size (normalized to average organoid size in control) of WT and tumor organoids on day 2 and day 3 post-seeding and treatment, p<0.0001 for WT and p=0.008 for tumoroids. C. Representative confocal immunofluorescence staining of YAP subcellular localization in WT organoids and tumor organoids after 60min treatment with MSN or control media. Overview images of whole organoids are maximum intensity projections showing three channels: YAP (magenta), Hoechst (blue), and β-catenin (yellow); scale bar = 50µm. Zoomed-in images represent single z-plane views of the areas highlighted in whole-organoid images, showing in order: merged channels, Hoechst channel only, and YAP channel only; scale bars = 50µm. Final zoomed images show close-up of the highlighted areas in the YAP channel; scale bars = 5µm. Dotted lines highlight the nuclei. D. Graphs show relative subcellular YAP localization (proportion of nuclear YAP to cytoplasmic YAP) in organoids when treated with MSN for 60min at day 2 post-seeding, compared with treatment with Control medium. Each point represents mean values from three biological replicates (n=3). For each biological replicate, a total of 5-7 images were analyzed to get mean values. Paired t-test, p=0.0015 and p=0.3971 for WT and tumor organoids, respectively. E. Graph shows the proportion of WT and Apc^Min/+^-derived tumor organoid cells expressing Sca-1 when exposed to MSN by flow cytometry. Points represent flow cytometry results from one organoid line (n=6 for WT and n=5 for tumor organoids). Paired t-tests, p=0.0256 and p=0.1503 for WT and tumor organoids, respectively. Error bars for all numerical graphs depict mean values +/-standard deviation.

We next investigated YAP activation in both WT and tumor organoids. Automated image segmentation of YAP immunofluorescence staining (Figure S1d) revealed a significant increase in YAP nuclear translocation, indicative of transcriptional activation, within 60 minutes following MSN exposure (Figure 1c,d). Tumor organoids showed constitutive YAP translocation independently of MSN exposure, characteristic of Apc^Min/+^ tumor intrinsic Wnt signaling activation (Figure 1c,d). YAP nuclear translocation was sustained at later time points, as observed 72h after exposure (Figure S2) and four days[22].

SCA1, a reparative and fetal-like stem cell marker known to be expressed downstream of YAP activation in the intestines[29], was analyzed using flow cytometry in MSN-treated organoids (Figure 1e and Figure S3). We found a significant expansion of SCA1^+^ cells in WT organoids when treated with MSN, from an average of 7.24% to 16.71%. In line with our results regarding YAP activation, untreated tumor organoids contained a significantly higher number of SCA1+ cells at baseline (average of 35.94%), which was not significantly different when exposed to MSN (average of 44.52%).

This data demonstrates that MSN exposure induces YAP activation in WT epithelial organoids and influences the growth of both WT and Apc^Min/+^-derived tumor epithelium.

### MSN elicits a regenerative-inflammatory phenotype in organoids, linked to Prostaglandin signaling and YAP/HIF1α activation

Besides its induction in intestinal tumors, YAP can be activated by various mechanisms, including mechanical cues induced by cell shape and cell tension[30], conditions experienced by organoids undergoing swelling. Therefore, to further investigate the molecular bases of the changes observed in MSN-exposed organoids, including the observed YAP activation, we analyzed early transcriptional responses to MSN treatment by bulk RNA-seq.

We characterized the gene signature of organoids after 20 min, 2 hours, or 24 hours of MSN exposure in healthy and tumor organoids, comparing their gene expression with that of control organoids not treated with MSN.

Early transcriptional changes showed no significantly differentially expressed genes (DEGs) at 20 min and only 2 upregulated DEGs after two hours of MSN exposure in healthy organoids (with a Benjamini-Hochberg false discovery rate of 20%) (Figure 2a). These corresponded to Ptger4 (EP4), a PGE2 receptor, and Nr4a1 (orphan nuclear receptor 4a1). Both genes are usually induced by inflammatory and stress-related cues and are heavily linked to prostaglandin signaling. Indeed, the most significant identified inducer of Ptger4 is PGE2[14]. The three other DEG hits at 2 hours on Figure 2a corresponded to three predicted pseudogenes of unknown function.

**Figure 2.**
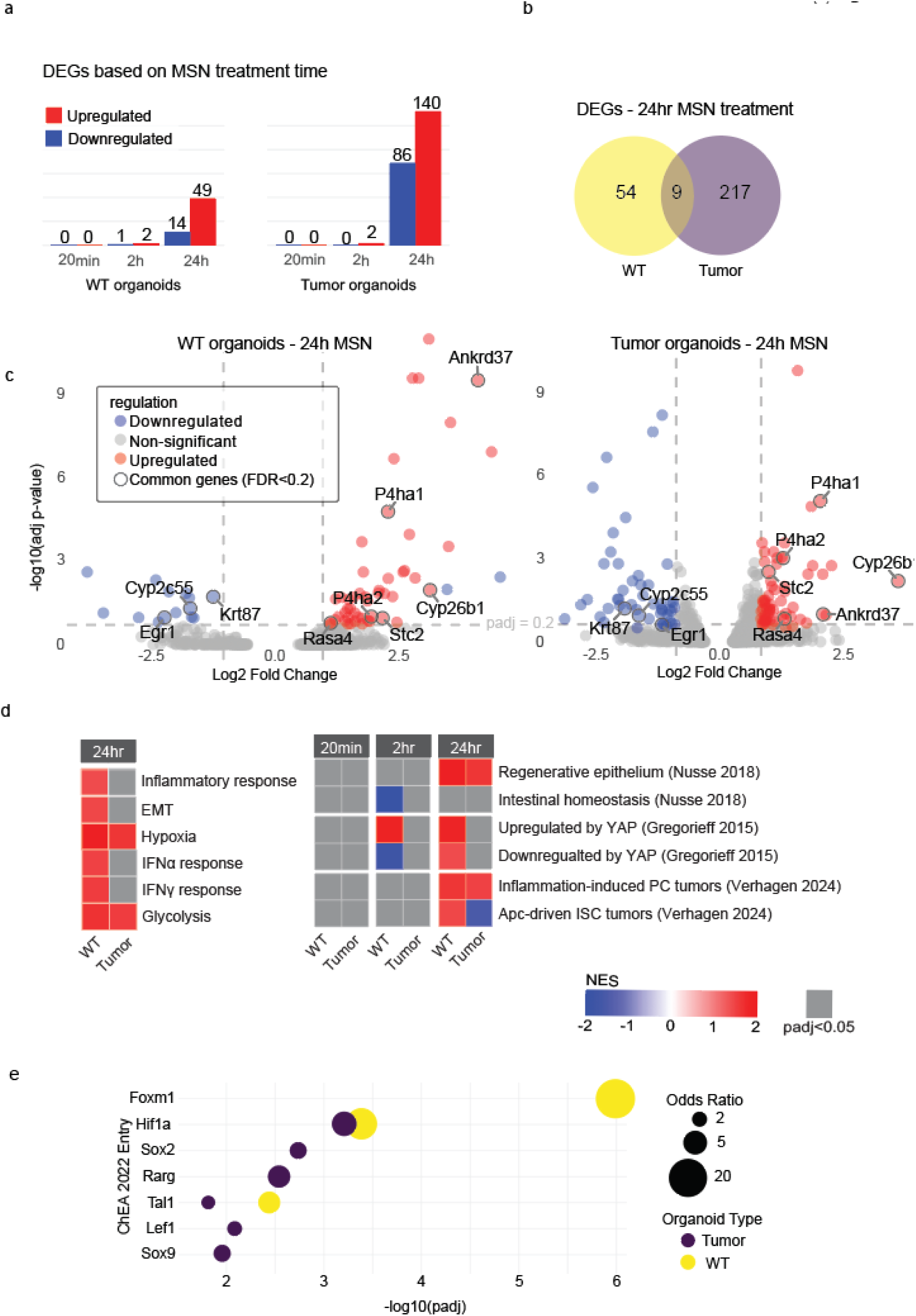
Characterization of gene expression responses to muscle-derived factors in WT and tumor organoids by bulk RNA-Seq. **A.** Bars indicate numbers of significantly differentially expressed genes (DEGs) in WT and tumor organoids treated with MSN for 20min, 2h, and 24h (Benjamini-Hochberg FDR<0.2). **B**. Venn diagram showing common and differentially expressed genes in WT and tumor organoids treated with MSN for 24h, compared to their respective organoids in control medium (Benjamini-Hochberg FDR<0.2). **C.** Volcano plots showing the distribution of differentially expressed genes in WT and tumor organoids treated with MSN for 24h, with commonly up-or downregulated genes between both WT and tumor organoids (Benjamini-Hochberg FDR<0.2) highlighted and labeled. **D.** GSEA on organoids treated with MSN for 20min, 2h and 24h, compared with untreated organoids. Normalized enrichment scores (NES) and significance for each gene set are shown on a color scale according to the provided legend. **E.** Gene overrepresentation analysis of differentially expressed genes at 24h after MSN treatment against the ChIP-X Enrichment Analysis Transcription Factor Targets 2022 (ChEA 2022) dataset. For all RNA-Seq data, n=3 for both WT and tumor organoids.

Most transcriptional differences, however, were observed after 24 hours of MSN exposure, with 63 and 226 DEGs, in WT and tumor organoids, respectively (Figure 2a-c). Among these 280 DEGs, we identified 9 that were commonly up-or downregulated in both sets of organoids (highlighted in the volcano plot). These shared genes likely represent a core transcriptional response to MSN at 24h, independently of the cellular context. Notably, four out of the six commonly upregulated genes were directly linked to Hypoxia-Inducible factor 1-alpha (Hif1α) activity, suggesting that MSN induces a conserved HIF1α signature across both WT and tumor organoids.

To gain further knowledge on the possible changes in biological processes induced by MSN exposure of epithelial organoids, we performed gene set enrichment analysis (GSEA) using hallmark gene sets from the Molecular Signatures Database (MSigDB) (Figure 2d). This analysis revealed that two related hallmark gene sets, hypoxia and glycolysis, were significantly enriched in both WT and tumor organoids after 24 hours of MSN treatment. Since YAP and HIF1α are described to interact and induce glycolysis, usually in cancer cells[31], it is not surprising to see here that the YAP (in WT organoids) and HIF1α activation (in both), induced by MSN, was associated with the glycolysis hallmark. Furthermore, we observed that MSN exposure elicited an inflammatory response in WT organoids, with enrichment in IFN-α and IFN-ϒ response hallmarks (Figure 2d).

GSEA using the YAP-target gene signature derived from Gregorieff et al. (2015)[28] showed a positive enrichment with YAP induction in WT organoids when exposed to MSN, while no significant enrichment was found in tumor organoids (Figure 2d), in line with our previous results on YAP activation (Figure 1). We also observed that MSN exposure induced an enrichment of “fetal-like” or regenerative signature in WT organoids (positive correlation with Nusse et al. 2018[32] gene set), also in line with our results on Figure1. In WT organoids exposed to MSN, when comparing with the Verhagen 2024 gene set[33], we observed an enrichment in tumoral features of APC-driven tumors as well as of tumors initiated by Paneth cells in response to inflammation. Tumor organoids, however, only showed an increase in features of inflammation-driven Paneth cell-derived tumors in response to MSN (Figure 2d).

Finally, we ran an overrepresentation analysis of DEGs in WT or tumor organoids after 24 hours of treatment with MSN using Enrichr[34] (Figure 2e). According to the ChIP-X Enrichment Analysis Transcription Factor Targets 2022 (ChEA 2022) dataset, targets of HIF1α were significantly overrepresented in both WT and tumor organoids in our gene expression dataset. In line, WT organoids treated for 24 hours with MSN showed significant enrichment of *Foxm1* targets, a cell-cycle regulator and proto-oncogene whose transcription is directly induced by HIF1α binding. Furthermore, among the transcription factors induced by MSN treatment, such as *Tal1* (common in WT and tumor organoids) and *Sox2, Rarg, Lef1,* and *Sox9* (in tumor organoids), are downstream targets of PI3K/Akt signaling (Figure 2d). Notably, it has been shown that YAP activates PI3K/Akt pathway[35], which acts upstream to stimulate the transcription, translation, and stabilization of HIF1α[36], suggesting a YAP-PI3K/Akt-HIF1α causative link, in WT organoids, and a sustained PI3K/Akt-HIF1α activation in tumoroids.

In sum, these data indicate that MSN exposure could be inducing an initial prostaglandin-mediated response resulting in sustained YAP and HIF1α stabilization/activation, and induction of an inflammatory, reparative state in WT organoids. Whereas HIF1α/hypoxia-glycolysis appears to be the dominant common response to MSN, YAP-dependent regenerative reprogramming is largely restricted to WT organoids.

### Smooth muscle-derived Prostaglandin E2 (PGE2) acts through Ptger4 (EP4) to induce MSN-driven phenotypical changes in organoids

Next, we sought to identify the muscle-derived molecule(s) responsible for the phenotypic switch-increased size and spheroidal morphology-observed in organoids exposed to MSN. To do so, we fractionated the components of our MSN by size, using gel-column separation (Figure 3a) and obtained 6 different fractions. The largest two fractions (Fractions 1 and 2) were expected to contain the majority of proteins (from 100kDa to 10kDa), and the following four fractions contained smaller molecules and ions and were virtually devoid of proteins (Fractions 3-6). We used these fractions to treat WT organoids and analyzed organoid growth using automated image analyses. As shown in Figure 3b-d, only two of the six fractions, fractions number 3 and 4, were able to induce the previously observed phenotypic switch toward larger organoids with a higher proportion of spheroids. These two fractions did not contain proteins but smaller molecules, indicating that the MSN-derived active compound, responsible for the observed morphological changes in organoids, was not a protein.

**Figure 3.**
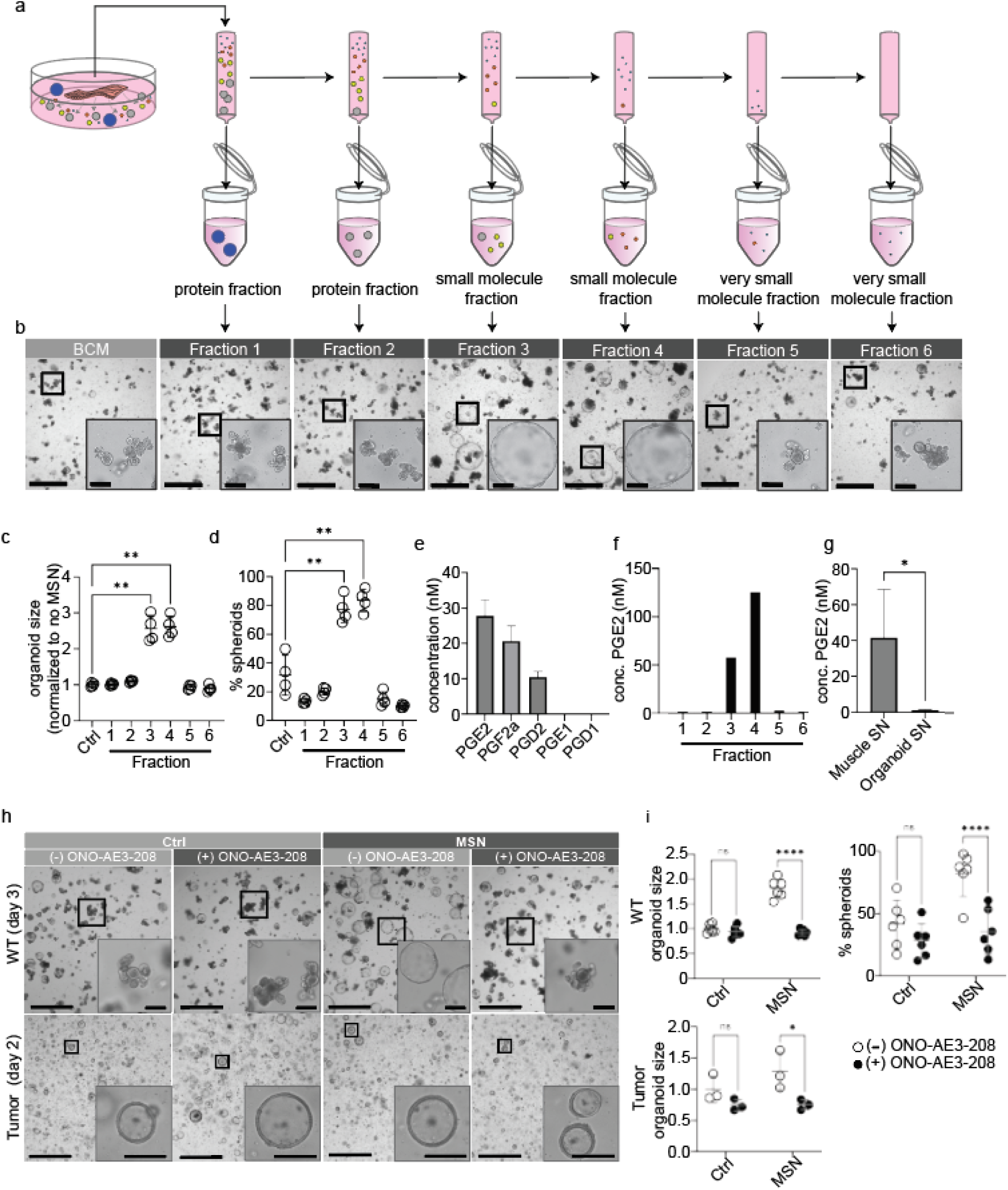
Identification of PGE2 as the MSN-derived driver of organoids’ phenotypical swift through EP4 binding. **A.** Schematic of MSN size column fractionation experiment. **B.** Representative bright-field images of WT organoids treated with size-fractionated MSN. Scale bars represent 1250µm and 200µm in zoomed insets. **C, D.** Graphs show organoid size and % of spheroids obtained through automated image quantification of WT organoids at day 4 post-seeding/treatment, where points represent mean values from four biological replicates (n=4), each treated with control (BCM) or the corresponding fractionated MSN. Data were analyzed using repeated measures one-way ANOVA. Asterisks show significant results from Dunnett’s multiple comparisons post hoc test. **C.** Relative organoid size of WT organoids treated with Control or size-separated MSN fractions, normalized to the mean organoid size of the control condition; p=0.0002, post-hoc padj= 0.079 and padj = 0.0029 for control vs Fraction 3 and control vs Fraction 4, respectively. **D.** Percentage of spheroids in each treatment group, p= 0.0001; post-hoc padj = 0.0025 and padj = 0.0046 for control vs Fraction 3 and control vs Fraction 4, respectively. **E**. Graph shows the concentration of different prostaglandins in nM, as determined by LC/MS (n=3 biological samples). **F**. Concentration of PGE2 in each MSN fraction as determined by PGE2 ELISA (n=1). **G**. Concentration of PGE2 from muscle supernatant compared to WT organoid supernatant in ELISA, n=10 and n=4, respectively. Unpaired t-test, p=0.0129. **H**. Representative bright-field images of WT organoids pre-treated or not with EP4 antagonist ONO-AE3-208, then treated with control (Ctrl) or MSN media. Scale bars represent 1250µm and 200µm for zoomed insets. **I**. Graph shows automated image quantification of size and percentage of spheroids in WT organoids and size of tumoroids, from the experiment described in (**H**) at day 3 and day 2 post-seeding, respectively. Points represent mean values from experiments on 6 organoid lines (n=6). Ṧídák’s multiple comparison test. For organoid size in WT: padj=0.5194, padj<0.0001 for control and MSN, respectively. For percentage of spheroids: padj=0.0536, padj=0.0003 for control and MSN, respectively. For tumor organoids, points represent mean values from experiments on 3 organoid lines (n=3). Ṧídák’s multiple comparison test, padj=0.1386, padj=0.0141 for control and MSN, respectively. All numerical graphs depict mean values +/-standard deviation.

Given this result, and in light of the bulk RNA-seq findings suggesting early activation of prostaglandin(PG)-related signaling in response to MSN, we next analyzed the content of PGs-small lipidic molecules-in MSN using LC-MS. As a result, we observed the presence of specific PGs in the MSN, with PGE2 being the most abundant, followed by PGF2a and PGD2 (Figure 3e). We next focused on PGE2 and analyzed its presence in the six size fractions of our MSN using ELISA. As shown in Figure 3f, we found PGE2 only in the two MSN size fractions that induced the organoids’ phenotypical switch, indicating that PGE2 was the small, non-protein molecule driving the phenotype.

We then compared the amount of PGE2 secreted by intestinal smooth muscle explants with the amount of PGE2 secreted by intestinal epithelial cells by measuring PGE2 concentration in MSN and medium conditioned by a comparable amount of organoid tissue (organoid supernatant (SN)) by ELISA. From this, we found that MSN medium contained 31 times higher levels of PGE2 than organoid conditioned medium (41.6 +/-8.5 nM in MSN versus 1.3 +/-0.1 nM in organoids SN) (Figure 3g), suggesting that the SMT is a major source of extracellular PGE2 in the intestines.

Previous literature has shown that exogenous treatment of PGE2, induces a similar spherical phenotype in WT organoids, linked to the activation of a reparative program in epithelial cells, promoting WAE - wound-associated epithelial cell-formation[10]. Furthermore, PGE2 downstream signaling has been shown to induce YAP nuclear translocation in epithelial cells[7, 13], which we had also observed in our organoids (Figure 1). In line with this, PGE2 signaling cascade induces PI3K/Akt activation, and increased PGE2 levels in epithelium are known to induce a regenerative fetal-like epithelial switch (YAP-dependent), inflammatory signaling, and HIF1α stabilization in the absence of hypoxia[10, 16], suggesting a causal link among the main MSN-induced transcriptional changes observed in our bulk RNA-seq (Figure 2).

We next treated organoids with exogenous PGE2 to see if it alone would cause the MSN-driven changes we observed in intestinal organoids. We found that treatment with either exogenous PGE2 or a stabilized form, 16,16-dimethyl PGE2 (dmPGE2), led to size and shape changes reminiscent of those caused by treatment with MSN (Figure S4a-d). As with MSN, WT organoids treated with PGE2 and dmPGE2 were significantly larger after MSN treatment (Figure S4a,b). Tumor organoids, also in keeping with our results from MSN treatment, showed an increase in size at day 2 post-treatment but lost any discernible differences by day 3 (Figure S4d). Both WT and tumor organoids treated with PGE2 had similar YAP activation patterns as observed when treated with MSN (Figure S4e, f). WT organoids also had higher proportions of SCA1+ cells following treatment with PGE2 or dmPGE2, while tumor organoids showed no significant differences (Figure S4g). In short, these experiments suggest that PGE2 is the main driver of MSN-observed phenotype on organoids.

PGE2 can act via different EP receptors (EP1 to EP4). The intestinal epithelium is known to express EP4 (*Ptger4*) in relation to WAE generation and tumor development[7]. We confirmed that the action of muscle-derived PGE2 is mediated through EP4 binding in our organoids by pre-treating organoids with ONO-AE3-208, a non-reversible EP4 inhibitor (Figure 3h,i). ONO-AE3-208 pre-treatment in WT organoids showed normal organoid size and morphology upon MSN exposure, suggesting that the morphological switch induced by MSN in organoids is led by PGE2-EP4 binding (Figure 3h,i). In tumor organoids, when treated with muscle supernatant, ONO-AE3-208 also inhibited the characteristic muscle supernatant-driven increase in size at day 2 (Figure 3h,i).

Taken together, our results indicate that PGE2 is the main SMT-derived component that, via PGE2-EP4-YAP axis, induces a spheroidal reparative/inflammatory phenotype in epithelial organoids.

### PGE2 is synthesized in the smooth muscle tissue (SMT) by transcellular biosynthesis

PGE2 synthesis is necessary for normal muscle homeostasis, mediating contraction or relaxation of the SMT depending on its distribution and specific receptor binding[37, 38]. Furthermore, its role in regulating regeneration after muscle injury (in skeletal muscle) by impacting muscle stem cells through EP4 binding has been recently described[39]. PGE2 synthesis pathway begins with cPLA2 (encoded by *Pla2g4a*), which hydrolyzes membrane phospholipids to release arachidonic acid. Then, COX1 and COX2 (encoded by *Ptgs1* and *Ptgs2*, respectively) convert arachidonic acid to PGH2, followed by conversion of PGH2 to PGE2 by one of three PGE2 synthases, PTGES, PTGES2, or PTGES3[40].

We reanalyzed bulk RNA-Seq data from our previous publication[22] to determine the expression of upstream components of PGE2 signaling in the intestinal tissue. By comparing the expression of PGE2 synthases between epithelial crypts and the SMT, we observed that the SMT expresses significantly higher levels of COX1 (*Ptgs1*) and COX2 (*Ptgs2*) than the epithelium (Figure 4a). PGE2-specific synthases *Ptges*, *Ptges2*, and *Ptges3*, were similarly expressed in epithelium and muscle. When we looked at PG receptors, however, we saw that the muscle expresses EP1-4 (*Ptger1-4*) while the epithelium predominantly and significantly expressed higher levels of EP4, followed by EP3 and 1, with negligible EP2 levels (Figure 4a).

**Figure 4.**
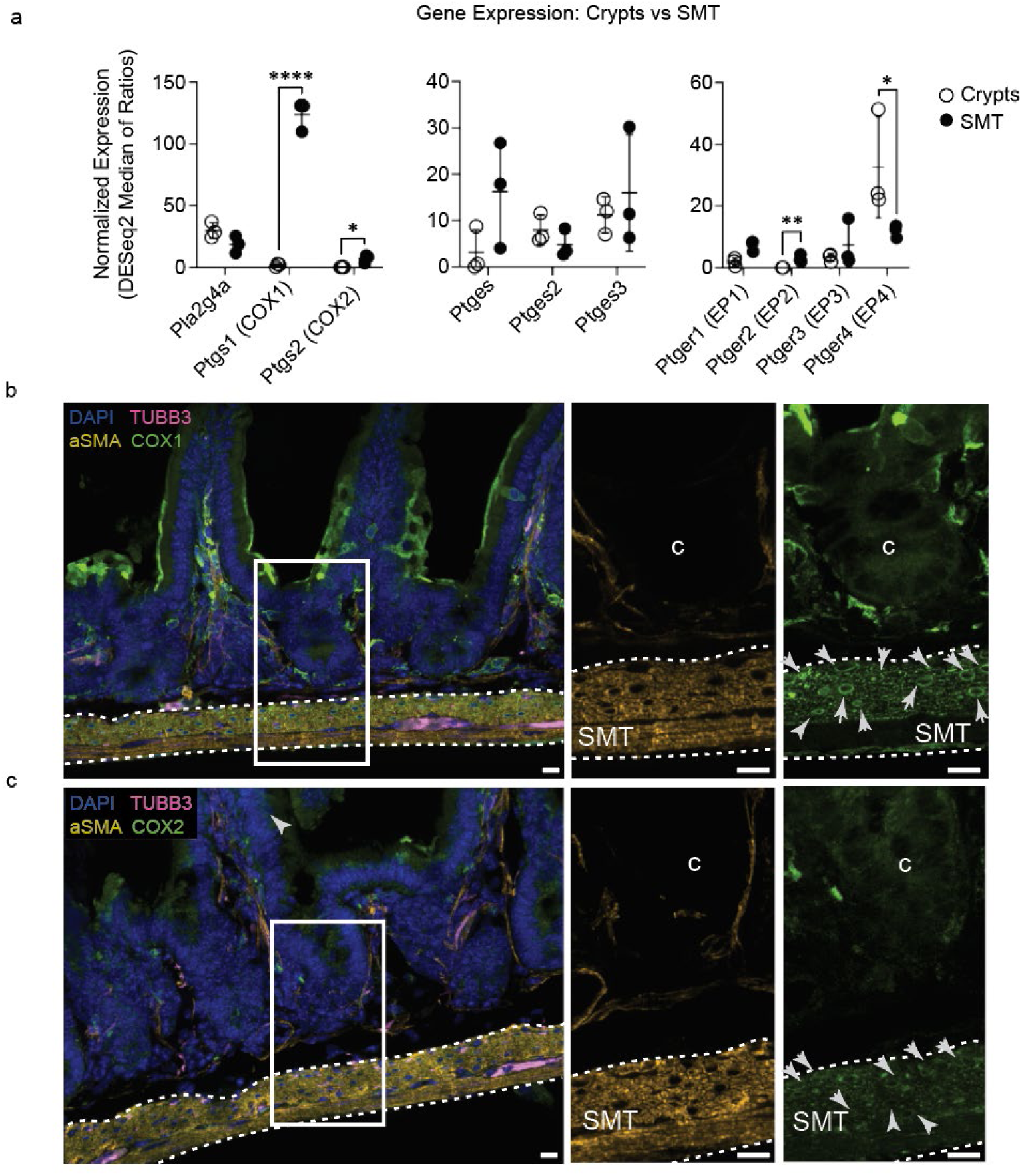
PGE2-converting enzymes expression and cellular distribution in the SMT. **A.** Graphs show bulk-RNAseq normalized gene expression of PGE2-converting enzymes and PGs receptors in epithelium (crypts) vs the SMT (n=3). **B.** Representative confocal immunostaining images of WT mouse intestinal sections showing COX1 staining (green) in crypts (C) and SMT. Notice the perinuclear membrane staining. The SMT is delineated by white dotted lines and stained for α-SMA (orange). The myenteric plexus is shown by TUBB3 (pink) and nuclei by DAPI (blue). Scale bar represents 10 µm. **C.** Representative confocal immunostaining of WT mouse intestinal sections showing COX2 staining (green) in crypts (C) and SMT, as in **B**.

To more closely examine the ability of the SMT to produce PGE2, we used fluorescent immunohistochemistry to co-stain for markers of resident cell types within the SMT, along with the PGE2 converting enzymes, in WT mice. In the SMT, COX1 and COX2 were expressed predominantly by α-SMA (alpha-smooth muscle actin)-positive smooth muscle cells (SMCs) located in the circular muscularis propria (Figure 4b,c). We also found positive COX1 expression together with immune CD45+ cells (Figure S6). Furthermore, we observed relevant COX1 and COX2 expression in the nervous plexus and serosa. In these two areas, COX1 and COX2 expression was limited to CD45+ cells and mesenchymal cells co-expressing MSLN and WT1 mesenchymal markers (Figure S6 and S7). Since both COX1 and 2 are membrane-binding enzymes, we observed the characteristic nuclear, endoplasmic reticulum, and, to a lesser extent, cytoplasmic membrane signal (Figure 4, S6 and S7).

We then analyzed the staining pattern of the enzymes directly upstream of PGE2 synthesis: PTGES, PTGES2, and PTGES3 (Figure S5a, c, e). We found that, although the three synthases show unique staining patterns, all PGE2 synthases showed areas of heightened expression that colocalize with Tubb3+ nerve cells, in close location with c-Kit+ cells marking interstitial cells of Cajal (ICC) (Figure S5a-f). Co-staining with smooth muscle cell marker α-SMA showed that these points of specific enrichment occur in areas negative for α-SMA+ smooth muscle myocytes (Figure S5g, h).

Transcellular biosynthesis has been described for PGE2 synthesis[41, 42], in which COX1/2-mediated initial synthesis steps occur in a defined cell type (e.g. α-SMA+ cells in the SMT), whereas the terminal conversion to PGE2 is carried out by a different cell population expressing PTGES, PTGES2 and/or PTGES3. In this case, our data indicate that the final PGE2 synthesis in healthy SMT occurs at neuro-muscular modulation areas and nervous plexuses within the muscle.

In sum, we found significant expression of PGE2-converting enzymes in the SMT in homeostasis.

### Tumor-associated muscle undergoes structural and molecular changes that include increased PGE2-converting enzymes expression

Given the prominent role of PGE2 in inducing a tumor-initiating gene expression program in the epithelium when expressed by stromal PDGFRa+ cells near the bottom of the epithelial crypt[7], we next examined potential changes in PGE2 expression in the SMT beneath the tumor (hereafter referred to as tumor-associated SMT) in Apc^Min/+^ mouse intestines.

When analyzing COX1 (*Ptgs1*), COX2 (*Ptgs2*), *Ptges, Ptges2* and *Ptges3,* and PGs receptors expression levels by bulk-RNAseq in tumor-associated SMT *versus* healthy (WT) SMT, we observed a significant increase in *Ptgs1* (COX1) expression in tumor-associated SMT, whereas the other converting enzymes and PGs-receptor remained similarly expressed, except for a significant decrease in *Ptger3* (EP3) receptor (Figure 5a). For methodological reasons, we could not compare healthy SMT and tumor-associated areas within the same mouse Apc^Min/+^ intestine in bulk RNA-seq, because tumors are widely distributed along the epithelium, and obtaining sufficient healthy SMT material would require using a different intestinal region. As intestinal regions differ in epithelial and mesenchymal specific features[43, 44], macroscopic regions of SMT underneath epithelial tumors were directly processed and compared to equivalent areas in the same anatomic regions in WT intestines. Later, using microscopy, we compared healthy SMT in tumor-distant areas and tumor-associated SMT within the same Apc^Min/+^ mouse. Importantly, no significant differences were observed or quantified when comparing the SMT in a WT mouse and the tumor-distant SMT in an Apc^Min/+^, in terms of structural changes or PGE2-converting enzymes staining patterns and levels (Figure 5 and S8).

**Figure 5.**
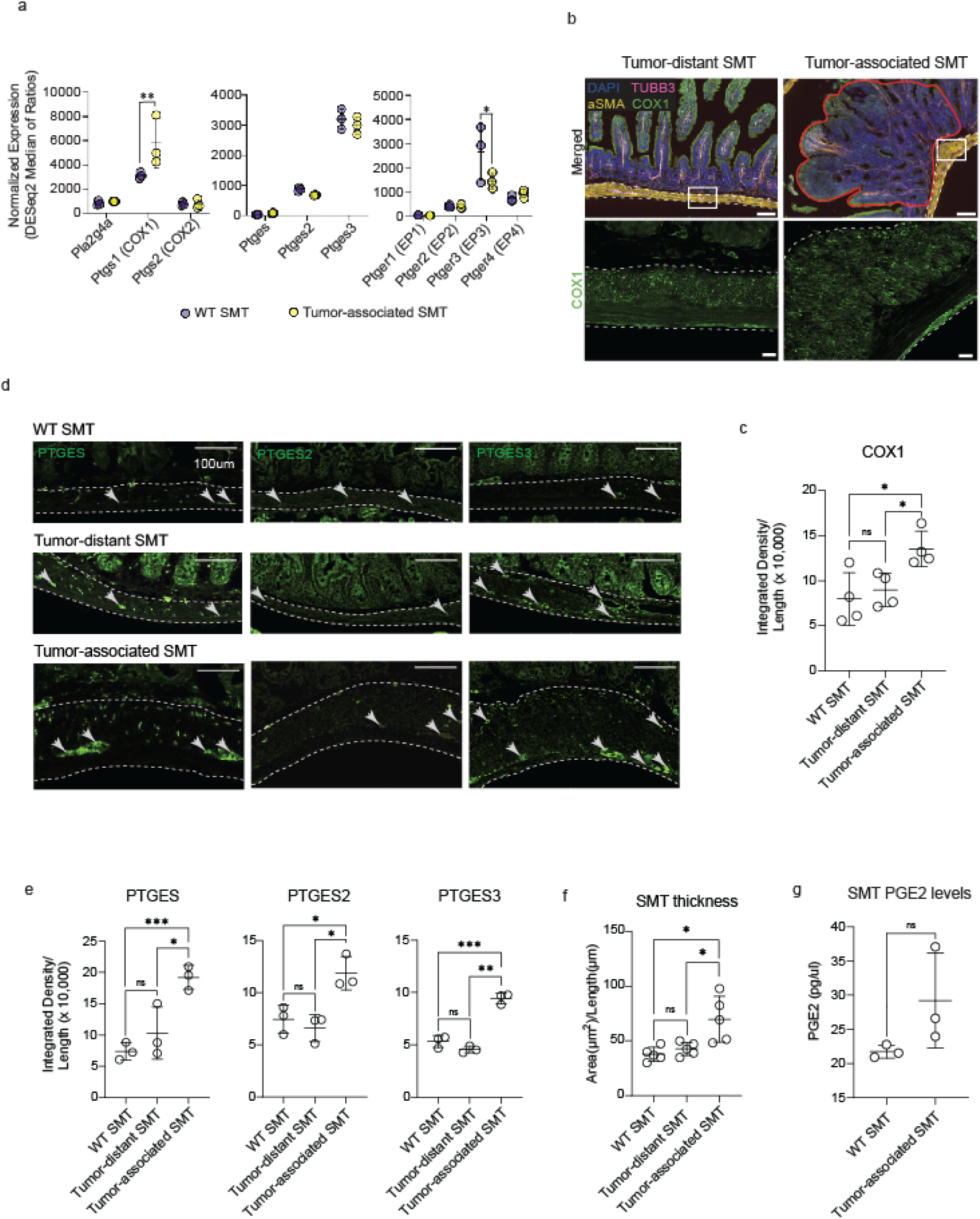
PGE2 signaling is increased in SMT under tumors. **A.** Differential gene expression of PGE synthesis pathway genes in healthy versus tumor-associated SMT. Negative binomial distribution; Benjamini-Hochberg adjustment FDR<0.001489 and FDR=0.027616 for Ptgs1 and Ptger3, respectively; and FDR>0.05 for others. Points represent normalized gene expression (DESeq normalization method) from one mouse, and error bars depict mean values +/-standard deviations. **B.** Representative confocal images of transversal sections of Apc^Min/+^ mouse intestine showing COX1 (Ptgs1) staining (green) in tumor-distant and tumor-associated SMT. The SMT is delineated by white dotted lines and stained for α-SMA (orange). The myenteric plexus is shown by TUBB3 (pink) and nuclei by DAPI (blue). Scale bars represent 100 µm in wide-view images and 10 µm in zoomed boxes. Tumoral epithelial area is highlighted by a red line. **C.** Graph shows integrated density measures of intensity of COX1 staining in intestinal smooth muscle of WT (WT SMT) and Apc^Min/+^mice (tumor-distant and tumor-associated SMT), normalized for muscle length. Each dot represents the mean results of immunohistochemistry intensity measurement from one mouse, using data from 3-5 images per mouse (n = 4). Data were analyzed with unpaired t-tests for comparing WT SMT vs tumor-distant SMT (p=0.5895) and WT SMT vs tumor-associated SMT (p=0.0198). Tumor-distant SMT and tumor-associated SMT were compared with paired t-test (p=0.0228). **D.** Representative confocal images of immunohistochemistry staining of mouse intestinal smooth muscle tissue, showing localization of PTGES, PTGES2 and PTGES3 in green in WT SMT, tumor-distant SMT and tumor-associated SMT. The SMT is delineated by white dotted lines. Scale bars represent 100µm. **E.** Graphs show integrated density values of PTGE synthases staining in intestinal smooth muscle under the mucosa of mouse small intestine, normalized for length. Each dot represents the mean results of immunohistochemistry intensity measurement from a mouse, using data from 3-5 images per mouse (n = 3). Statistical analyses were done as for C. with unpaired t-tests to compare WT and Apc^Min/+^ SMT and with paired t-tests to compare tumor-distant and tumor-associated SMT. PTGES intensity; WT SMT vs tumor-distant SMT; p=0.3150; WT SMT vs tumor-associated SMT; p=0.0010; and tumor-distant SMT vs tumor-associated SMT; p=0.02080. PTGES2 intensity; WT SMT vs tumor-distant SMT; p=0.4843; WT SMT vs tumor-associated SMT; p=0.0212 and tumor-distant SMT vs tumor-adjacent SMT; p=0.0239. PTGES3 intensity; WT SMT vs tumor-distant SMT; p=0.1129; WT SMT vs tumor-associated SMT; p=0.0007 and tumor-distant SMT vs tumor-adjacent SMT; p=0.0096. **F**. Graph shows quantification of SMT thickness in WT (WT SMT) and Apc^Min/+^ mice (tumor-distant and tumor-associated SMT). Each dot represents the mean muscle area from one mouse (n=5), with measurements in Apc^Min/+^ mice taken either in areas under a tumor or in non-tumoral regions, where each point shows average data from 3-11 images. Statistical analyses were done as for C. and E. with unpaired t-tests to compare WT and Apc^Min/+^ SMT (WT SMT vs tumor-distant SMT; p=0.2545; WT SMT vs tumor-associated SMT; p=0.0113) and with paired t-tests to compare tumor-distant and tumor-associated SMT (p=0.0179). **G**. Graph shows PGE2 levels in healthy vs tumor-associated SMT measured by LC/MS (n=3 biological samples). Mann-Whitney test; p=0.1. All numerical graphs depict mean values +/-standard deviation.

We next analyzed COX1 and COX2 protein expression patterns comparing healthy (or tumor-distant) SMT and tumor-associated SMT in Apc^Min/+^ intestines (Figure 5b-g). Like in healthy SMT (Figure 4), PGE2 synthases pattern confirmed the expression of COX1/2 enzymes in the SMCs of the tumor-associated SMT, predominantly located in the circular muscularis propria and mesenchymal cells at the serosa of mouse intestines (Figure 5, Figure S6, and S7), whereas PTGES, PTGES2, and PTGES3 were mostly expressed by αSMA-negative cells in tumor-associated SMT (Figure S8b).

COX1 and COX2 staining showed an expansion of the positive area for both enzymes in tumor-associated SMT (Figure 5b and S6 and S7 for COX2 staining). When signal integrated density was quantified (a measure of the total fluorescence signal in a given area), we observed significantly higher COX1 levels within the tumor-associated SMT, in line with our bulk-RNAseq data (Figure 4a).

PTGES, PTGES2, and PTGES3 staining comparing WT SMT, tumor-distant, and tumor-associated SMT in Apc^Min/+^ intestines showed significantly increased integrated fluorescence density in tumor-associated SMT, when normalized for intestinal length (Figure 5d,e).

To note, we consistently observed a significant SMT structural alteration with an increase in the thickness of the tumor-associated SMT, under epithelial adenomas (Figure 5f). Indeed, the difference in PGE2 synthases staining disappears when normalized for the area of the muscle measured, instead of the length (Figure S8a). These results indicate an expansion of the SMT compartment underneath the adenoma, with larger numbers of cells expressing PGE2 converting enzymes. We also observed an increased CD45+ infiltrate within tumor-associated SMT, as shown in Figure S6, possibly related to the inflammatory signature of the tumor-associated SMT found in our bulk-RNAseq. These infiltrated cells were generally negative for COX1/2 staining (Figure S6).

COX1 and COX2 expression are often correlated with PGE2 levels in tissues[45]. Furthermore, the increased thickness of the tumor-associated SMT expressing PGE2 converting enzymes might indicate increased production and availability of the lipid in these areas. To analyze if the observed transcriptional changes and protein expression profile will result in locally increased PGE2 levels, we performed LC/MS analysis of PGE2 in SMT lysates from healthy and tumor-associated SMT, finding a trend towards increased PGE2 levels in tumor-associated SMT (Figure 5g).

Taken together, these data indicate that PGE2 synthesis is increased within the tumor-associated SMT primarily through transcriptional upregulation of *Ptgs1* (COX1), likely leading to elevated levels of the pro-tumorigenic, inflammatory lipid PGE2 beneath the tumor.

### Tumor-associated SMT shows increased inflammation and decreased contractile markers expression

We next decided to comprehensively characterize the changes affecting this tumor-associated SMT by performing bulk RNA-Seq comparing healthy SMT and tumor-associated SMT from WT and Apc^Min/+^ mice, respectively. Among the most significantly upregulated genes in tumor-associated SMT (see volcano plot on Figure 6a), we found genes related to ECM remodeling, erythropoiesis and hemoglobin synthesis and metabolism, and markers of inflammation.

**Figure 6.**
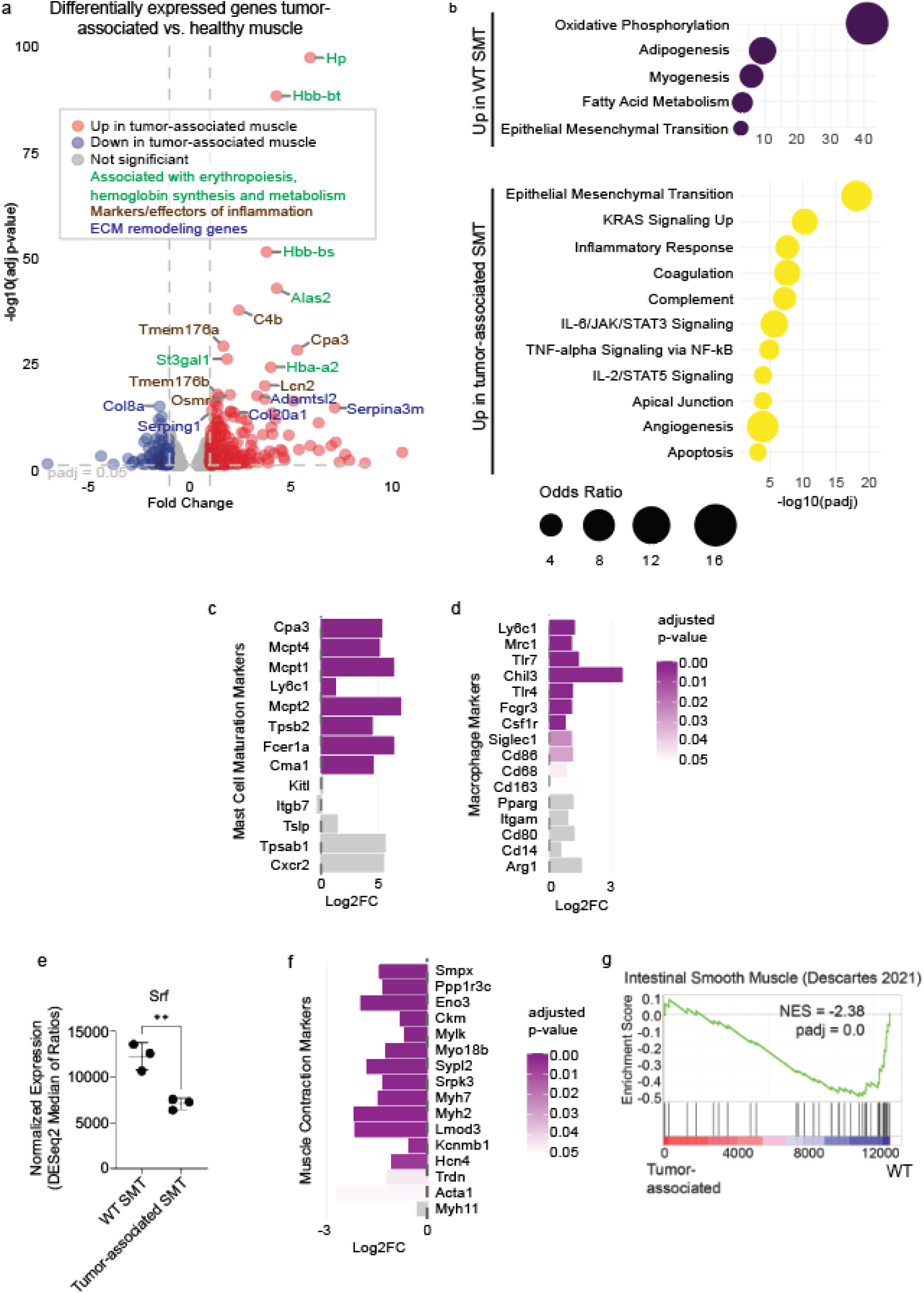
Tumor-associated SMT has increased expression of inflammation and immune cell markers, and decreased smooth muscle contractile markers. **A.** Volcano plot shows differentially expressed genes in tumor-associated muscle compared to WT muscle. The top-most differentially expressed genes (by Benjamini-Hochberg corrected p-value) are highlighted. Genes significantly upregulated in tumor-associated muscle are marked in red, and genes significantly downregulated in tumor-associated muscle are highlighted in blue. **B**. Graphs show overrepresentation analysis of genes upregulated and downregulated in tumor-associated muscle. Hallmarks for which genes are significantly overrepresented in tumor-associated muscle are shown in yellow, and hallmarks for which genes are significantly overrepresented in WT muscle are shown in purple. Size of the dots reflects odds ratio, and the position of the dots reflects the adjusted p-value of the overrepresentation. **C, D**. Gene expression changes in mast cell maturation markers and macrophage genes, respectively, in tumor-associated muscle compared to WT muscle. The direction of the bars depicts the change in gene expression as measured by log2 fold change, and the color of the bars indicates the adjusted p-value as corrected by Benjamini-Hochberg adjustment. **E.** Expression of the transcription factor *Srf* in WT healthy muscle compared to Apc^Min/+^ tumor-associated muscle, Negative binomial distribution; Benjamini-Hochberg adjustment FDR<0.0001. Points represent normalized gene expression (DESeq normalization method) and error bars depict mean values +/-standard deviations. **F**. Gene expression changes in SRF targets. The direction of the bars depicts the change in gene expression as measured by log2 fold change, and the color of the bars indicates the adjusted p-value as corrected by Benjamini-Hochberg adjustment. **G.** GSEA of tumor-associated muscle versus WT muscle against a gene set of intestinal smooth muscle during development. For all bulk RNA-Seq data, n=3 for both tumor-associated muscle and healthy muscle.

Using Enrichr, we checked gene expression changes for overrepresentation of the Hallmarks from MSigDB. Among the 289 significantly upregulated genes in tumor-associated SMT, we found upregulation in pathways related to inflammation, including hallmarks of Inflammatory response, complement activation, IL-6/JAK/STAT3 signaling, TNF-α signaling via NF-kβ, and IL2-STAT5 signaling. We also found enrichment for angiogenesis. Interestingly, tumor-associated SMT has an increased gene signature related to EMT and KRAS signaling, two pathways usually related to tumor regions, and particularly found in CRC tumors (Figure 6b).

Furthermore, we found that many of the most upregulated genes in tumor-associated SMT were markers of mast cell maturation, but not activation or migration (Figure 6c), and upregulation of macrophage markers in tumor-associated SMT over WT healthy muscle (Figure 6d). Interestingly, overrepresentation analysis of downregulated genes in tumor-associated SMT showed signaling related to oxidative phosphorylation, adipogenesis, and myogenesis (Figure 6b). In line with this, we found a muscle-specific decrease in *Srf* expression, a key transcription factor regulating muscle development and function[46–48] (Figure 6e), and whose transcription and function can be repressed by PGE2[18]. Tumor-associated SMT showed a significant reduction in genes related to SMT function, in particular, a significant downregulation in muscle contractile markers such as *Ckm, Mylk*, *Kcnmb1,* and *Myh11*, suggesting an alteration in SMC homeostasis and differentiation (Figure 6f). GSEA analyses showed a significant negative Normalized Enrichment Score (NES) in the hallmark of Intestinal Smooth muscle, indicative of a significant downregulation in muscle genes (Figure 6g).

Taken together, these results show that the SMT is heavily affected by the tumoral epithelium prior to carcinoma progression, potentially weakening the muscle by reducing contractile gene expression and increasing inflammation within the SMT. The observed PGE2 increased levels in tumor-associated SMT could contribute to the increased inflammation, immune cell recruitment, and SMC dysfunction found in these muscles. However, it will be of special interest to identify the tumor-derived molecules driving the observed changes in tumor-associated SMT.

### Identification of tumor-secreted proteins and their possible impact on the SMT

The shifts found in tumor-associated SMT suggest alterations in normal epithelial-SMT crosstalk, in which signaling cues originating from the tumoral epithelium may drive the observed modifications in the SMT.

In accordance with our bulk RNA-seq data comparing WT and Apc^Min/+^ derived tumor organoids, among the tumor organoid signature (1703 genes significantly upregulated in Apc^Min/+^ tumor organoids), we found enrichment of genes related to biological processes such as cell migration, epithelial-to-mesenchymal transition (EMT), and blood vessel morphogenesis (Figure 7a). Among the most enriched pathways, we found TGFβ signaling, which, in established tumors, is known to actively reprogram the tumor microenvironment, promoting metastasis, angiogenesis, EMT, matrix remodeling, and immune suppression[49]. Considering that TGFβ is a key regulator of the SMT as well, the altered TGFβ signaling induced by epithelial tumors could have an impact on SMT fitness[49].

**Figure 7.**
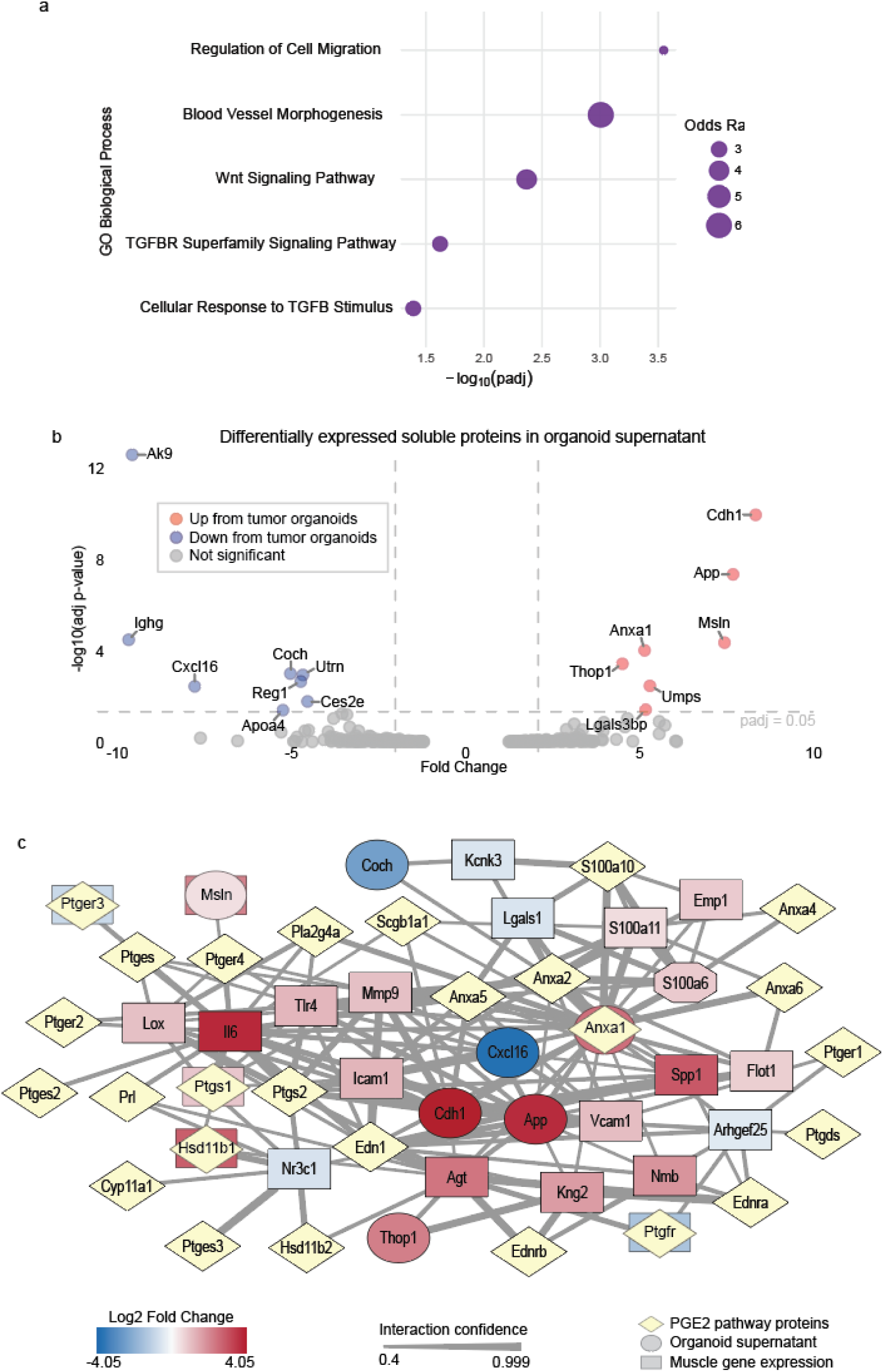
Construction of protein interaction network proposes possible novel mechanisms of epithelium-to-muscle mediated crosstalk. **A.** GO Biological Process overrepresentation analysis showing enriched terms among genes upregulated in tumor organoids. B. Volcano plot showing the differentially expressed soluble proteins secreted by tumor organoids compared to WT organoids. Proteins occurring more frequently in tumor organoids supernatants are shown in red, while proteins occurring more frequently in WT organoids supernatants are shown in blue. Proteins with significantly differential protein expression between tumor and WT organoids are identified and labeled. For the proteomic experiment, n=4 for both WT and tumor organoids supernatants. C. A gene interaction network between proteins with the biggest changes in expression in organoid supernatant with or without a tumor (oval), genes that are significantly up-or downregulated in tumor-associated muscle (rectangle), and the PGE2 pathway proteins that may affect PGE2 production in the muscle (yellow diamond). The log2 fold change in protein expression from organoid supernatant proteomics or in gene expression from muscle bulk RNA-Seq are shown by a color scale, as shown in the legend. Edges shown here depict interactions that connect organoid supernatant factors to PGE2 pathway proteins, either directly or through one differentially expressed gene in the muscle. Edge thickness depicts the total interaction score for each interaction as documented by the STRING database.

To identify the possible molecules directly secreted by the tumoral epithelium that could be implicated in this tumor-SMT crosstalk, we performed quantitative proteomic analysis on the supernatant (SN) obtained from Apc^Min/+^ tumor organoids and compared it with the supernatant derived from healthy WT organoids. Of the soluble proteins released by tumoroids, we identified seven proteins significantly more abundant and eight proteins significantly less abundant in tumor organoids supernatant compared to WT organoids supernatant (Figure 7b). Among the upregulated proteins in tumoroids SN, we found proteins typically highly expressed by tumoral cells in CRC, such as Annexin1 (ANXA1), Galectin-3-binding protein (LGALS3BP), mesothelin (MSLN), and Amyloid precursor protein (APP), linked to different cancer progression processes, including cell proliferation, migration, aggressiveness, poor prognosis, and fibroblast activation. However, when analyzing the reported impact of these proteins in the SMT, we observed their described implication in SMT fitness, suggesting a possible tumor-SMT way of communication, which could explain some of the shown tumor-associated features in the SMT. For example, LGALS3BP expression has been related to a phenotypical switch in SMCs to an inflammatory synthetic phenotype in vascular smooth muscle[50], MSLN activates fibroblasts and contributes to tumor progression, affecting the tumor microenvironment[51], and APP leads to vascular SMCs degeneration when accumulated in the brain, with putative similar effects in other regions SMCs[52]. Interestingly, all these upregulated proteins have been linked to TGFβ.

We investigated whether the differentially secreted factors in tumor organoids were able to affect gene expression in their associated muscle counterparts, particularly the possible interaction with the PGE2 pathway. To do this, we used the STRING database[53] to build an interaction network, linking the differentially expressed soluble factors derived from tumor and WT organoids to PGE2 synthesis-related proteins, either directly or through intermediate genes that we identified as differentially regulated in the muscle in tumor-associated compared to healthy WT SMT. With this, we highlighted interaction pathways by which intestinal tumors could drive the observed changes in tumor-associated SMT (Figure 7c, Table S1).

In sum, our data suggest the presence of active communication between the tumoral epithelium and the underlying SMT, resulting in structural, cellular, and molecular significant changes in the SMT. Such changes include alterations in PGs signaling, with an upregulation of COX1, and its downstream product PGE2, and a reduction in SMT differentiation and increased inflammation. The impact of such changes in muscle performance or their implications in tumor cell migration through the SMT and metastasis needs further investigation.

## Discussion

In this study, we reveal a bidirectional communication between the epithelium (healthy and tumoral) and the underlying SMT. We characterize this communication in the Apc^Min/+^ mouse model, which usually develops non-invasive adenomas, suggesting an early onset of this bidirectional communication. By using organoid culture and SMT-conditioned medium, we demonstrated that SMT-derived factors, such as PGE2, can lead to phenotypic changes in epithelial organoids reminiscent of a tumor-like state. Additionally, we showed that the presence of an epithelial tumor remodels the underlying SMT, inducing a gene expression program that increases expression of PGE2 production machinery and lowers expression of muscle master regulator SRF, along with muscle contraction genes, and inflammation. Such SMT-induced changes would create a more permissive environment for the tumoral cells to grow unrestricted, due to the known PGE2 pro-tumoral effect, and simultaneously weaken the natural tumoral boundary, the SMT.

Previous work from our group[22] and other groups[23, 24] has demonstrated that the SMT is a source of growth factors, such as BMP antagonists, known to regulate the epithelial SCs niche. Here, we demonstrate that the SMT is also a source of the pleiotropic lipid mediator, PGE2. PGE2 plays a central role in SMT physiology, contributing to the regulation of muscle tone and muscle stem-cell fitness[20, 21]. Beyond the SMT, PGE2 is involved in proinflammatory responses and is also essential for intestinal repair. Thus, tight regulation of PGE2 production and signaling is critical in intestinal physiology.

In 2017, Miyoshi et al[10] demonstrated that treatment of intestinal organoids with exogenous PGE2 induces a reparative epithelial program (WAE), characterized by nuclear accumulation of β-catenin, independently of canonical Wnt activation, needed for epithelial healing after injury. Besides showing the same morphological changes in organoids when exposed to MSN, our transcriptomics data confirmed the activation of a reparative signature in the epithelium that includes some similarities to Apc-driven tumors and inflammation. We demonstrated that such changes were governed by the presence of PGE2 in the MSN, resulting in a rapid phenotypic shift characterized by increased YAP activation/HIF1α signaling and the expansion of a subpopulation of reparative fetal-like stem cells (SCA1+ cells) in WT organoids.

Notably, the MSN-induced phenotypic shift in organoids, including YAP activation and Sca1+ cell expansion, was markedly less pronounced in tumor organoids. These findings suggest that the role of PGE2 in tumor development may lie in promoting tumor initiation rather than enhancing later tumor aggressiveness. Consistent with this interpretation, Roulis et al. (2020) showed that ablation of PGE2 production in stromal *Ptgs2*-expressing fibroblasts located adjacent to epithelial crypts significantly impaired intestinal tumor formation, highlighting the functional requirement of stromal-derived PGE2 for tumor initiation[7]. Although the SMT is not directly apposed to intestinal crypts, changes in this tissue, including alterations in PGE2 levels, could impact other intermediate cells within the mucosa, and given that PGE2 usually elicits a self-amplifying positive loop, local levels could be amplified, ultimately reaching the epithelium. Moreover, the extensive remodeling observed in tumor-associated SMT is likely to influence tumor formation and progression through mechanisms extending beyond PGE2 signaling alone. For example, we detected an upregulation of genes related to erythropoiesis and hemoglobin synthesis and metabolism. The presence of extramedullary hematopoiesis due to hypoxia and stress has been previously described, as well as the tumor-induced erythropoiesis, supporting our idea of this established two-way communication between the SMT and the tumor.

Previously, we sought to identify the molecules responsible for the phenotype shift observed in MSN-treated organoid cultures. In 2021, we identified the SMT-secreted protein Periostin, which, when added to organoid cultures, induced a modest increase in organoid size[22]. However, Periostin alone did not fully recapitulate the morphological changes induced by MSN. At the time, we did not investigate the contribution of non-protein components present in the MSN. Our current findings identify PGE2 as the principal driver of the phenotype; nevertheless, additional factors within the MSN may still contribute to the modulation of organoid behavior. Indeed, the combined activity of multiple MSN-derived factors may act synergistically to regulate organoid growth and differentiation. Consistent with this, although PGE2 was the most abundant PG within the MSN, we also identified the presence of additional PGs that may also contribute to epithelial regulation in homeostasis or in the presence of a tumor.

We characterize the structural and molecular alterations found in the tumor-associated SMT, using the Apc^Min/+^ mouse model. Our data suggest that the tumoral epithelium is able to remodel the underlying SMT, inducing a gene expression program that increases expression of *Ptgs1*, a necessary enzyme in the PGE2 synthesis pathway, and decreases expression of *Srf*, a master transcriptional regulator of muscle development, proliferation, and differentiation[46–48]. The simultaneous observation of these two changes is not surprising, as PGE2 has been shown to have an inhibitory effect on *Srf* transcription[18]. The downregulation of this gene could explain some of the other observations in tumor-associated SMT, in particular, the downregulation of muscle-related contractile machinery, since SRF is a master regulator of contractile genes. Indeed, knockout of SRF has been shown to lead to dilation of the intestinal tract and impaired contraction of intestinal smooth muscle[46, 54]. So, in conditions such as CRC, a local increase in PGE2 production could be an important driver in processes such as immune evasion and tumor growth stimulation. Whether SMT-associated alterations in PGE2 signaling contribute to epithelial tumor progression remains unknown.

We propose that this local increase in PGE2 signaling may be responsible for other observed changes in the tumor-associated SMT. PGE2 can induce macrophage migration and infiltration into inflamed tissue, and activated macrophages (via COX2-induction mainly) can be a major source of PGE2[55]. On the other hand, PGE2 is a known chemoattractant for mast cells[56], able to modulate mast cell function, including these cells’ maturation, activation, and cytokine release in a receptor-dependent and context-specific manner[57]. Furthermore, increased mast cells in tumor-associated SMT could contribute to the breakdown and weakening of tumor-associated SMT through degranulation and release of its effector molecules, processes known to negatively impact SMT fitness[58–62]. Therefore, the observed immunological shift in the tumor-associated SMT can be, at least partially, reflective of the described increased PGE2 production in these regions.

Notably, we observed that the different PGE2-converting enzymes were expressed by different cellular types in the SMT. Initial steps of PGE2 synthesis will occur in α-SMA+ cells (principally expressed by mature smooth muscle cells within the smooth muscle wall) that express relatively high levels of *Ptgs1* (COX1), while terminal PGE2 synthases will complete the synthesis in a neighboring cell expressing Ptges, Ptges2, or Ptges3 enzymes. This mechanism of transcellular biosynthesis has been previously demonstrated for PGE2 synthesis, allowing to enhance or bypass of the limited biosynthetic capacity of a single cell type, especially in cases of high demand, such as inflammation, injury response, or during tumoral growth [41, 42].

For decades, non-steroidal anti-inflammatory drugs (NSAIDs), including COX1/2 inhibitors, have been known to protect against CRC, partly by downregulating PGE2 levels in the intestine. However, broad inhibition of prostaglandin synthesis showed unwanted side effects, producing gastrointestinal problems, because of the fundamental role of PGEs in intestinal (mucosal and muscular) homeostasis[63]. These observations are consistent with our findings, which highlight the essential role of prostaglandins, in particular PGE2, in intestinal homeostasis, especially within the SMT, and the importance of having a tight regulation of PGE2 (and PGs) among the different intestinal compartments.

In this work, we also proposed several candidate crosstalk mechanisms by which tumors can enact paracrine signaling to the underlying SMT. Of note, we found that the chemokine CXCL16, which plays a role in the maintenance of muscle cell adhesion, proliferation, and regeneration[64, 65], is released at significantly lower levels from tumor epithelial organoids than from healthy small intestinal organoids. We also identified lowered amounts of extracellular UTRN, the intracellular counterpart of which plays a role in the maintenance of muscle function[66]. We also found higher levels of a soluble ANXA1 and APP in our tumor-associated SMT. A soluble form of ANXA1 has been shown elsewhere to promote pro-inflammatory pathways in neutrophils, alluding to the fact that soluble ANXA1 could be responsible for an increased inflammation profile in tumor-associated SMT. Finally, soluble fragments of APP, sAPPα and sAPPβ, have been shown to decrease cellular adhesion in other cell types[67], and thus may also work to weaken the SMT.

Looking ahead, several avenues of investigation could clarify the functional relevance of the pathways we identified. Directly testing our proposed mechanisms, for example, by treating healthy SMT with recombinant versions of the candidate factors we identified or by selectively blocking others using neutralizing antibodies, would help determine whether these signals are sufficient or necessary to drive the transcriptional changes we observed in tumor-associated SMT. Moreover, our research establishes a proof-of-concept of the importance of considering intestinal SMT in the study of intestinal tumor development. An important next step, therefore, will be to assess whether SMT remodeling has functional consequences for disease progression.

Determining whether the smooth muscle alterations we observed can influence tumor growth dynamics, SMT invasion, or clinical outcomes is needed and could be crucial for shaping future therapeutic strategies in intestinal cancer.

## Materials and Methods

### Mice

Apc^Min/+^ mice in the C57BL/6 background (C57BL/6J-ApcMin, stock no. 002020) were obtained from The Jackson Laboratory. All mice were housed and maintained at the Comparative Medicine Core Facility at NTNU (CoMed). Mice were handled under pathogen-free conditions and mouse breeding and experiments were performed following Norwegian legislation on animal protection and were ethically approved by the local governmental animal care committee. Apc^Min/+^ breeding and experiments were approved in advance by the Norwegian Food Safety Authority (FOTS IDs: 17072 and 30058).

Experiments on WT and Apc^Min/+^ mice were performed at 3-4 months of age to allow sufficient time for spontaneous tumors to form and were carefully monitored to avoid situations of moderate to high pain. End point protocols were applied when needed, following ethical procedures laid out by the CoMed Facility at NTNU. All mice were genotyped by PCR from ear punch tissue with the following primers: fw: 5’-ttccactttggcataaggc-3’; ApcWT-rev: 5’-gccatcccttcacgttag-3’; Apc^Min^-rev: 5’-ttctgagaaagacagaagtta-3’. Euthanasia of mice was performed by induction with isoflurane followed by cervical dislocation.

### Small intestine crypt isolation

Small intestinal crypt isolation was performed following a modified protocol, originally published by Sato et al [68]. In brief, 10 cm of proximal small intestine was extracted, and the lumen was rinsed with ice-cold, cell culture-grade PBS (Sigma-Aldrich D8537). The intestine was then cleaned of surrounding fatty tissue and cut open longitudinally. Then, the villi and mucus were removed by gently scrapping the intestinal luminal side with a coverslip. The intestine was then cut into small pieces of 3-4 mm in length using a scalpel blade, and tissue pieces were rinsed repeatedly in ice-cold PBS by shaking gently in a 50mL conical vial, removing the debris-laden PBS, and adding clean PBS, and repeating until the PBS rinse was clear. Then, the tissue pieces were incubated in 2mM EDTA in PBS at 4°C with gentle rocking for 30 min. Following incubation, the crypts were dissociated by transferring to a new 50mL conical vial with ice-cold PBS and shaking vigorously. The tissue pieces were allowed to settle at the bottom of the conical vial and the dissociated crypts, in suspension in PBS, were passed through a 70µm strainer and collected in a tube, both pre-coated with 50-100% Fetal Calf Serum (FCS). This step was repeated several times to collect a sufficient number of crypts. The suspension was then centrifuged at 300 x g, 5 min, at 4°C, and resuspended in basal culture medium (BCM; Advanced DMEM-F12 (Gibco 12634-010), PenStrep (Sigma-Aldrich P0781), 10 mM HEPES (Gibco 15630-056), 2 µM GlutaMAX (Gibco 35050-061)) to get the appropriate density of crypts for seeding.

### Wild-type intestinal organoid culture

Crypts were seeded at a density of 250-300 crypts per 40-50 µL droplet containing 70% Matrigel (Corning 356231) and 30% BCM. The droplets were allowed to polymerize at 37°C for 5 min in pre-heated 24-well plates, then submerged in medium containing basal culture medium, 20% R-Spondin 1 conditioned medium (gift from Calvin Kuo), 10% Noggin conditioned medium (gift from H. Clevers), 1X B27 (Gibco 17504001), 1X N2 (Gibco 17502001), 1mM N-acetyl-L-cysteine (Sigma-Aldrich A7250), and 50ng/mL mouse recombinant EGF (Gibco PMG8041). Organoids were cultured at 37°C and 5% CO_2_. Culture medium was replaced every 2-3 days until passaging was required.

Organoids were passaged at day 4-6 post-seeding by disrupting the Matrigel droplets with 1000 mL pipette tip. The organoids and polymerized Matrigel were further mechanically disrupted with repeated passages through a syringe and blunt-tipped needle (BD 303129) against the wall of a 50 mL conical vial on ice. The suspension was centrifuged at 300 x g, 5 min, at 4°C and resuspended in a fresh mixture of 70% Matrigel and 30% BCM before seeding in new droplets at a 1:3 dilution. If confocal imaging was the experimental endpoint, the organoids were seeded in 8-well chamber slides (ibidi 80801), at a density of ∼100 crypts or organoid fragments per 30 µL droplet containing 70% Matrigel and 30% BCM instead of normal 24-well plates. As with seeded organoids, passaged organoids were given fresh medium every 2-3 days until further passaging was required.

Following experimental treatments, organoids were monitored using live imaging daily using the EVOS FL Auto 2 live imaging system.

### Apc^Min/+^-derived tumor organoid seeding and culture

Intestinal tumor organoids were obtained from the small intestine of a 3–4-month-old Apc^Min/+^ mouse. Following euthanasia, the small intestine was removed, and the lumen was rinsed with ice-cold, cell culture-grade PBS, then cut open longitudinally and surveyed for tumors. Five to ten tumors were excised from the first 10 cm of the proximal end of the small intestine with a round-edged scalpel. The tumors were then finely minced with a scalpel and washed with ice-cold, cell culture-grade PBS by gently shaking in a 50 mL conical vial and removing the liquid once the tissue pieces had settled at the bottom. The tumor mince was incubated in 2mM EDTA for 30 min at 4°C with gentle rocking. Following this, the EDTA was removed, and the tumor mince was incubated with 0.5 mg/mL collagenase at 37°C for 20 min with intermittent agitation. The collagenase was then removed, and the tumor mince was further dissociated by passing it through a blunt-tipped needle against the wall of a 50 mL conical vial. The dissociated tumoral tissue was then passed through a 70 µM cell strainer and collected in a conical vial, both pre-coated with 50-100% FCS. The cell suspension was serially diluted to 1:10 and 1:100 with cell culture-grade PBS and centrifuged to pellet cells. Each dilution, including an undiluted sample (1:1), was resuspended in 70% Matrigel and 30% basal culture medium and seeded as 40-50 µL droplets as previously mentioned. The Matrigel and cell mix was allowed to polymerize for 5 minutes at 37°C on a pre-heated cell culture plate, then covered with tumor organoid medium consisting of BCM, 1X B27 (Gibco 17504001), 1X N2 (Gibco 17502001), 1mM N-acetyl-L-cysteine (Sigma-Aldrich A7250), and 50 ng/mL mouse recombinant EGF (Gibco PMG8041) to select for tumor cell growth. The medium was replaced every 2-3 days until passaging was required.

### Intestinal smooth muscle isolation and generation of MSN

Intestinal smooth muscle tissue (SMT) samples were obtained from the first 10 cm of mouse small intestine. The intestinal lumen was flushed with ice-cold cell culture-grade PBS to remove food and kept submerged in ice-cold PBS. The outside of the duodenum was cleaned of connective tissue and fat under a dissecting microscope, cut open longitudinally, and rearranged such that the luminal side was facing down. A shallow cut was made in the muscle tissue, and the muscle tissue was peeled back gently with a sharp, rounded scalpel blade.

To generate MSN, the muscle tissue was kept on ice in cell culture-grade PBS until ready to weigh, gently dabbed dry on a lint-free absorbent towel, and weighed on a microscale. The muscle was then rinsed twice in 40 mL of sterile 0.9% saline and placed a volume of BCM proportional to the weight of the muscle (1mL for 20 mg of muscle). Supernatants were collected 12-24 hours following incubation.

To compare tumor-associated and healthy epithelium-associated smooth muscle, the entire small intestine was removed from a 3–4-month-old Apc^Min/+^ or wild-type mouse. The distal end (2-9 last cm) of the small intestine was removed. The intestines were cleaned of connective tissue and fat under a dissecting microscope, cut open longitudinally, and inspected for tumors. Apc^Min/+^ mice of this age often have this section covered in tumors, and the sample was discarded if too few tumors were present. The intestines were then arranged such that the luminal side was facing down and cut into 10-14 pieces of equal size (5-7 mm long). The muscle was peeled off each tissue piece using a rounded scalpel blade. The smooth muscle pieces were stored in RNAlater (Invitrogen AM7020) or RNAprotect Tissue Reagent (Qiagen 76104) for RNA extraction, or stored in ice-cold cell culture-grade PBS for subsequent RNA extraction or supernatant generation following the previously mentioned procedure.

### Treatment of organoids with MSN, MSN fractions, PGE2, or exogenous PGE2

Immediately after passaging, organoids were submerged in normal organoid medium, MSN-supplemented medium (wherein the basal culture medium is replaced by a 1:1 mixture of basal culture medium and MSN, MSN), normal organoid medium supplemented with 2.9 nM PGE2 (Cayman Chemical 14010, 5 mg/mL primary stock dissolved in DMSO), or 2.9 nM dmPGE2 (R&D Systems 4027, 10 mg/mL in methyl acetate). The organoids were then allowed to grow at 37°C and 5% CO2 without additional media changes until the experimental endpoint, at which point they were harvested for downstream analysis.

For shorter experiments where seeded organoids would not have time to develop in the treatment time, for example in the 60 min treatment condition in the YAP internalization experiment and the bulk RNA-Seq experiment, organoids were first seeded in normal media and allowed to grow for two days. The media were then replaced with either normal organoid medium or MSN-supplemented medium.

### Treatment of organoids with Ptger4 inhibitor, ONO-AE3-208

Immediately upon seeding, organoids were pre-treated with media supplemented with 10 uM of ONO-AE3-208 (Cayman Chemical 14552-1, 10 mM primary stock dissolved in DMSO) for 60 min. Following this, the media was replaced with either normal organoid medium or tumor organoid medium, either supplemented with MSN or not.

### Gel column fractionation of MSN

An initial amount of 2 mL of MSN was fractionated by gel filtration using PBS at a flow rate of 0.5 ml/min using Superdex 75 10/300 at 4°C. The initial 30 fractions obtained were then pooled based on elution time into 6 main fractions, then aliquoted and frozen at-80°C for PGE2 ELISA or organoid treatment (see section above).

### Quantification of PGE2 and other prostaglandins in conditioned media

Quantification of PGE2 in MSN was performed using a PGE2 ELISA (Thermo Fisher Scientific, EHPGE2) following instructions provided by the manufacturer. The plate was read with a Bio-Rad iMark microplate reader, and the standard curve was generated with a 5-parameter fit with Bio-Rad Microplate Manager.

Quantification of PGE2, PGD1, PGD2, PGE1, and PGF2α in normal MSN, as well as comparison of PGE2 levels in tumor-associated muscle compared to healthy muscle, were performed using LC-MS at the Lipidomics Core Facility, Danish Cancer Institute. To prepare the sample, 150 µL of MSN was mixed with 15 µL of an internal standard mixture containing 5 µM of each deuterated prostaglandin (PGD1-d4, PGD2-d4, PGE1-d4, PGE2-d4 and PGF2α-d4), corresponding to 75 pmol of each internal standard per sample. Following this, 900 µL ethyl acetate was added and the samples were shaken at 2000 rpm for 5 min at 4 °C, then centrifuged at 6000 × g for 5 min. The organic phase was then transferred to a new tube and evaporated to dryness. The remaining residue was dissolved in 60 µL acetonitrile/water (20:80, v/v) and mixed for 1 min, followed by centrifugation at 6000 × g for 5 min to remove sediment. A volume of 50 µL of the supernatant was transferred to LC–MS vials for analysis.

### Liquid chromatography

Chromatographic separation was performed on an UltiMate 3000 micro HPLC system (Thermo Fisher Scientific) equipped with a 0.5 × 150 mm YMC-Triart C18 ExRS analytical column with 0.5 μm particle (YMC, Kyoto, Japan). Mobile phase A was LC–MS grade water, and mobile phase B was LC–MS grade acetonitrile containing 10 mM ammonium acetate. The flow rate was 20 µL/min, the column was maintained at 35 °C, and the injection volume was 20 µL. The gradient was programmed as follows: 0– 0.5 min, 20% B; 0.5–0.6 min, 30% B; 0.6–8.8 min, 50% B; 8.8–8.9 min, 50% B; 8.9–9.9 min, 75% B; 9.9– 10 min, 20% B; and 10–11.5 min, 20% B for column re-equilibration.

### Mass spectrometry for PGs identification

The LC system was coupled to a Q Exactive quadrupole-Orbitrap mass spectrometer (Thermo Fisher Scientific). Samples were analyzed in negative ion mode using a full scan (MS) range of m/z 300–500. The instrument was operated at a resolution of 35,000 (at m/z 200) with one microscan per scan. The maximum injection time was 100 ms, and the automatic gain control (AGC) target was 1 × 10^5.

For validation and confirmation in parallel to full MS, MS/MS spectra were acquired using targeted parallel reaction monitoring (tPRM) for four m/z values (m/z 351.2177, 353.2334, 355.2428, and 357.2584). These m/z values cover both endogenous prostaglandins and their corresponding deuterated internal standards. The tPRM scans were acquired at a resolution of 17,500 (at m/z 200) with an AGC target of 5 × 10^5, a maximum injection time of 128 ms, one microscan, a quadrupole isolation window of 1.1 units, and a normalized collision energy (nCE) of 27.5%.

### Immunofluorescence staining of organoids

Organoids were seeded at 30-40 organoids per 30 µL droplet (70% Matrigel, 30% basal culture medium) in 8-well chamber slides (Ibidi 80801). Organoids were either immediately treated with medium supplemented with MSN, external PGE2, or control medium and allowed to grow for three days; or grown for two days to allow for partial differentiation to occur and treated for 60 minutes. After each experiment, media was removed from the wells and rinsed three times with PBS. Organoids were fixed with 4% paraformaldehyde and 2% sucrose solution in PBS at RT for 30 min. The fixing solution was removed by very gentle pipetting and the wells were rinsed very carefully with PBS three more times. The slides were then stored at 4°C for staining.

To immunostain, organoids were permeabilized and blocked in 100mM glycine, 5% normal goat serum, and 0.3% Tx-100 in PBS for 30 min at RT. The wells were then rinsed twice with PBS, allowing each wash to incubate without agitation for 5 min at RT. The wells were then incubated with primary antibodies against YAP (1:100, Cell Signaling Technology 14074S), and beta-catenin (1:200, BD Biosciences 610154) overnight at 4°C with gentle agitation. The wells were then washed three times with PBS-Tween 0.1%, allowing for 5 min incubation at RT each time, in gentle agitation. Secondary antibodies AlexaFluor 555 (1:500, Invitrogen A21422) and AlexaFluor 647 (1:500, Invitrogen A21245), along with 1:1000 Hoechst dye was incubated overnight at 4°C with gentle agitation. Then, the wells were washed three times with PBS-Tween 0.1% as described before, then rinsed once with distilled water for 5 minutes. The wells were then covered in 150 µL of Fluoromount G (Invitrogen 00-4958-02) before imaging.

### Organoids imaging, quantification, and automated image analyses

#### Instance segmentation of intestinal organoids and spheroids

Bright-field images of live organoid cultures were acquired using an EVOS FL Auto 2 live imaging system and a LPlan-Achromat 2X/0.06NA air objective (EVOS AMEP 4631). z-stacks per well were processed plate-wise using a custom pipeline written in Python[69] (https://doi.org/10.5281/zenodo.20085163). Files were grouped by well_id extracted from EVOS-compatible filenames, excluding non-z-stack images. For each well, z-planes were collapsed by minimum-intensity projection and saved as one projection image per well. Instance segmentation was performed on each projection with a pretrained YOLOv8 model [70] (https://doi.org/10.5281/zenodo.20118755) (./models/GPU_1px_cle_eroded_labels.pt) at confidence threshold 0.6, with classes differentiated (organoids), undifferentiated (spheroids), and dead. Predicted instance masks were parsed per class and quantified with skimage.measure.regionprops to extract area, filled area, perimeter, circularity, eccentricity, and solidity for each object. Per-object measurements were exported as plate-level CSV files, then aggregated to well-level summaries by computing class counts, mean morphology descriptors, and derived ratios (dead_ratio, organoid_ratio, spheroid_ratio).

#### YAP Nucleus to Cytoplasm ratio quantification

Fluorescence images of fixed organoids were obtained on a Zeiss LSM880 Airyscan using a Plan-Apochromat 20X/0.8NA/0.55 WD (420650-9902) air objective. Fluorescently immunostained organoid 3D stacks (.czi) were analyzed with a CellposeSAM[71]-based nuclei segmentation workflow[72] (https://doi.org/10.5281/zenodo.20081226) followed by morphology-guided cytoplasm simulation. For each image, voxel scaling metadata was extracted to compute anisotropy-corrected nuclei inference in 3D. Predicted nuclei labels were filtered by volume to remove outliers, and per-nucleus cytoplasmic shells were generated by controlled dilation. Nuclei were mapped to parent organoid masks to preserve organoid context. YAP signal quantification was performed as mean intensity in nucleus and simulated cytoplasm for each label, together with nucleus/cytoplasm volumes. A per-label YAP nuclear-to-cytoplasmic ratio (nuclei_cyto_ratio) was then computed. Batch outputs were consolidated into CSV files, filtered using volume thresholds and orphan-label exclusion (cells not belonging to an organoid), and summarized at technical-replicate and biological-replicate levels.

### Flow cytometry

Matrigel droplets containing organoids were disrupted using a sterile P1000 pipette tip or cell scraper and transferred along with media into a 15 mL conical vial. The contents were centrifuged at 300 x g, 4°C for 5 min to pellet the organoids. 1 mL of TrypLE (Gibco 12603-010) was added to each vial and incubated at 37°C for 30 min, triturating with a P1000 pipette every 10 min to prevent clumping cells. Cells were then centrifuged at 500 x g, 4°C for 3 min to remove the TrypLE and resuspended in PBS with 2% FCS. AlexaFluor 488-conjugated Sca-1 antibody (1:400, BioLegend, 108116) was used as the surface stain and DAPI (1:1000) was used to identify non-viable cells. Dyes were incubated for 30 min on ice. The cells were washed twice in PBS + 2% FCS by centrifugation and resuspension in fresh buffer. The flow cytometer used was BD FACSCanto II. FlowJo 10.10 was used for downstream analysis of flow cytometry data.

### RNA Extraction from organoids and muscle tissue

RNA was isolated from organoids using the *Quick*-RNA Microprep kit (Zymo Research, R1050/1) and from smooth muscle tissue using the RNEasy Plus Mini Kit (Qiagen, 74034), following instructions from the manufacturers. Followinoid experiments, 3-4 Matrigel droplets containing organoids were collected and broken down by multiple passages through a blunt-tipped needle against the wall of a 50 mL conical vial on ice. The slurry was centrifuged at 300 x g, 4°C for 5 min, then the supernatant was removed. 1mL lysis buffer from the *Quick*-RNA Microprep kit was added to the pellet, and the remaining RNA isolation procedure was carried out in accordance with the manufacturer’s instructions. For smooth muscle RNA isolation, tissues were submerged in RNAlater (Invitrogen AM7020) or RNAprotect Tissue Reagent (Qiagen 76104) and kept at 4°C or-20°C until RNA extraction. Tissues were then placed in a homogenization tube (Precellys P000912-LYSK-A) with 500uL cell lysis buffer with 5uL β-mercaptoethanol. Samples were then lysed using the Precellys 24 Homogenizer (Bertin Technologies) for 4 cycles of 15s at 6500 rpm, with 5 min of rest on ice between the second and third homogenization cycles. The samples were centrifuged at 10,000 x g and 4°C for 2 min, to remove the foam, then transferred to a clean tube. The samples were then centrifuged again at 10,000 x g and 4°C for 5 min to remove the debris pellet. Following this, RNA extraction was carried out in accordance with the manufacturer’s instructions. RNA concentration and quality were determined by spectrophotometry using a DeNovix DS-11 Plus or a Nanodrop ND1000 spectrophotometer. RNA samples were only used for downstream experiments if they exceeded a concentration of 20ng/µL, total RNA over 200ng, with OD260/280 and OD260/230 ratios both over 2.0.

### Bulk RNA-Seq of organoids and muscle tissue

For bulk RNA-Seq of organoids, WT organoids and tumor organoids were seeded as described above in 24-well plates and grown in fully supplemented organoid medium. Media were replaced with MSN-supplemented organoid medium, or normal organoid medium for the control group, at day two post seeding for all time points, and the contents of each experimental group (3-4 Matrigel droplets each for Control, 20min, 2h, and 24h groups) were harvested for downstream RNA extraction at their respective time points after treatment with MSN-supplemented organoid medium. The Control, 20min, and 2h groups were harvested on day two, whereas the 24h group was harvested on day three post-seeding. The experiment was run separately on three different organoid lines for both WT and tumor organoids to generate three biological replicates for each.

For bulk RNA-Seq of muscle tissue, healthy and tumor-associated muscle were obtained from three WT mice for healthy muscle and three Apc^Min/+^ mice for tumor-associated muscle, using the technique described in the section: Intestinal smooth muscle isolation and generation of MSN.

Library preparation and sequencing were performed by Novogene UK. Messenger RNA was purified from total RNA using poly-T oligo-attached magnetic beads. After fragmentation, the first strand cDNA was synthesized using random hexamer primers, followed by the second strand cDNA synthesis using dTTP. The samples were then A-tailed, adapter ligated, size selected, amplified, and purified before sequencing using Illumina NovaSeq 6000. Mapping was done using hisat2, quantification was done using featureCounts, and differential gene expression analysis was done using DESeq2.

### Gene-set enrichment analysis (GSEA) and overrepresentation analysis

The GSEA was run with log2fold change as weights using the R package fgsea. Hallmark gene sets were obtained from MSigDB[73]. The gene set for inflammatory proliferation was obtained from bulk transcriptomic data generated by Nusse et al (2018), of mouse duodenal crypts infected with *Heligmosomoides polygyrus* or not [32]. The gene set of inflammation-induced Paneth cell tumors compared to Apc mutation-driven tumors was derived from bulk transcriptomic data from Verhagen et al., of tumors obtained from mice with induced Apc KO in Paneth cells and given water with dextran sulfate sodium (DSS), compared to tumors that spontaneously arose from mice with induced Apc KO in intestinal stem cells [33]. The gene set for Apc KO organoids and WT organoids were derived from data by Neerven et al [74]. The gene set that is up-and down-regulated by YAP were obtained from data by Gregorieff et al [28]. Intestinal cell type signatures were obtained from Haber et al [75]. Macrophage markers were generated manually from literature review. Finally, the intestinal smooth muscle development gene set was obtained from the Descartes gene expression during development project [76]. For overrepresentation analysis, a list of genes with significant differential expression (BH-adjusted p-value < 0.05) was constructed for WT organoids 24h post-treatment versus WT organoid control, as well as tumor organoids 24h post-treatment versus tumor organoid control. The gene lists were analyzed with the Enrichr browser application [77, 78].

### Generation of organoid conditioned medium from WT or tumor organoids

WT and tumor organoids were both seeded with a target density of 250 organoids per dome and immersed with organoid medium containing BCM, 20% R-Spondin 1 conditioned medium, 10% Noggin conditioned medium, 1X B27 (Gibco 17504001), 1X N2 (Gibco 17502001), 1mM N-acetyl-L-cysteine (Sigma-Aldrich A7250), and 50ng/mL mouse recombinant EGF (Gibco PMG8041), at 0.5mL per 50uL Matrigel dome. On day 2, the WT and tumor organoids were checked to see if they formed a comparable number of viable organoids. The medium was then replaced on day two and four after seeding. The conditioned medium from day two to four was then collected in Eppendorf tubes and flash-frozen in liquid N2 before storage in-80°C and thawed before downstream analysis or experiments.

### Mass spectrometry for conditioned media

Proteins were precipitated using methanol/chloroform from 0.2 ml conditioned growth media. The protein pellet was dissolved in 100µl 1% SDC, 100mM Tris-HCl pH-8.5, 10mM TCEP, 40mM CAA and heated at 85°C for 45 min. Each sample was added 100 µl 0.1M of Ammonium bicarbonate, 1 µg trypsin, and digested over-night at 37°C. Peptides were desalted using C18 spin columns, dried in a speedvac centrifuge and resuspended in 0.1% Formic acid before MS analysis. LC-MS/MS were performed on a timsTOF Pro (Bruker Daltonics) connected to a nanoElute (Bruker Daltonics) HPLC. Peptide separation was done using a Pepsep 25 (150um*25cm) column with running buffers A (0.1% formic acid) and B (0.1% formic acid in acetonitrile) with a gradient from 2% B to 40%B for 100min. The timsTof instrument was operated in the DDA PASEF mode with 10 PASEF scans per acquisition cycle and accumulation and ramp times of 100 ms each. The ‘target value’ was set to 20,000 and dynamic exclusion was activated and set to 0.4 min. The quadrupole isolation width was set to 2 Th for m/z < 700 and 3 Th for m/z > 800.

### Mass Spectrometry data analysis for conditioned media

Mass spectrometry raw data analysis Open workflow (Ref: PTM-Shepherd: Analysis and Summarization of Post-Translational and Chemical Modifications From Open Search Results - ScienceDirect) was used to inspect the raw data to determine optimal search parameters for MaxQuant version 2.6.5.0 with the integrated Andromeda search engine (Ref: MaxQuant enables high peptide identification rates, individualized p.p.b.-range mass accuracies and proteome-wide protein quantification | Nature Biotechnology). Raw files acquired on a Bruker timsTOF instrument in TIMS-DDA/PASEF mode were searched against the mouse reference proteome (taxonomy ID: 10090) and additional protein entries downloaded from UniProt together with MaxQuant’s internal contaminants database. Carbamidomethylation of cysteine was specified as a fixed modification, while oxidation of methionine, protein N-terminal acetylation, and deamidation of asparagine/glutamine were included as variable modifications. Enzyme specificity was set to Trypsin/P allowing up to two missed cleavages. The minimal peptide length was set to 7 amino acids and the maximal peptide mass to 4600 Da.

First-search peptide tolerance was set to 20 ppm and the main-search tolerance to 10 ppm. Label-free quantification (LFQ) and iBAQ quantification were enabled with a minimum ratio count of 1 peptide. Match-between-runs (MBR) functionality was enabled to reduce missing values between samples using a matching time window of 1 minute and an alignment window of 20 minutes together with ion mobility matching (0.05 Vs/cm² matching window and 1 Vs/cm² alignment window). If a peptide feature was transferred across runs without direct MS/MS sequencing, it was annotated as “By matching” rather than “By MS/MS” in the MaxQuant output tables.

Differential protein expression analysis was performed using the R package DEP using LFQ intensity values of identified proteins. Proteins or peptides identified as potential contaminants or reverse hits were removed prior to analysis. The threshold for missing values was 50% and imputation of missing values used the “MinProb”, q=0.01 method. LFC cutoff = 0.5 and significance cutoff = 0.1. Visualization of differential protein expression was performed using the R package, ggplot2.

### Immunohistochemistry and immunofluorescence analysis

Tissues were immersed in 4% formaldehyde (9713, VWR Chemicals) at 4°C for 12 hours for fixation, followed by 15% and 30% sucrose in PBS at 4°C until they sank for cryoprotection. They were then embedded in O.C.T. compound (361603E, CTL Chemicals) and stored at-80°C.

The embedded tissues were cut to 20-µm sections on Thermo Scientific CryoStar NX70 cryostat for immunofluorescent labelling. Heat-induced epitope retrieval was performed to all sections with pH-6 citrate buffer at 99°C for 15 minutes. The sections were blocked with 10% normal goat serum (S-1000-20, Vector Laboratories) or 10% normal donkey serum (ab7475, Abcam) in TBS with 0.3% Triton™ X-100 (1.08603 Millipore) for 1 hour at room temperature. They were then incubated in primary antibodies dilution, containing 1% respective serum, 1% bovine serum albumin (A7906 Sigma-Aldrich), 0.3% Triton™ X-100, 0.01% sodium azide in TBS, at 4°C for 18-48 hours. They were incubated in secondary antibody dilution, containing 0.3% Triton™ X-100 in TBS, for 1 hour at room temperature and counterstained with Hoechst 33342 (1:1000). The following primary antibodies were used: anti-PTGES rabbit polyclonal (1:50, 160140, Cayman Chemical), anti-PTGES2 rabbit polyclonal (1:25, 10881-1-AP, Proteintech), anti-PTGES3 rabbit polyclonal (1:50, HPA038672, Sigma-Aldrich), anti-CD45 rat monoclonal (1:100, 550539, BD Pharmingen), anti-TUBB3 mouse monoclonal (1:500, ab78078, Abcam), anti-c-KIT rabbit monoclonal (1:100, 3074S, Cell Signaling Technology), anti-PDGFR-alpha goat polyclonal (1:50, AF1062, R&D Systems), anti-MSLN mouse monoclonal (1:100, ab236546, Abcam), anti-PTGS1 rabbit monoclonal (1:100, ab109025, Abcam), anti-PTGS2 rabbit monoclonal (1:100, 12282T, Cell Signaling Technology). Cy3™-conjugated anti-α-smooth-muscle-actin mouse monoclonal (1:200, Sigma-C6198, Aldrich), Alexa-Fluor-647-conjugated anti-WT1 rabbit monoclonal (1:100, ab283325, Abcam). The secondary antibodies with corresponding host and target animals were Invitrogen’s Alexa-Fluor-dye-conjugated antibodies. In the case where the primary-antibody host species were the same, Tyramide SuperBoost™ kits were used for immunofluorescence labelling (Alexa Fluor™ 488 and 647 tyramides, Invitrogen) according to the manufacturer’s instructions.

Labelled-tissue images were acquired on a Nikon TI2 confocal microscope with a Crest V3 spinning disk unit connected to a Teledyne Photometrix Kinetix sCMOS camera; the objectives used were Plan Apochromat λD 20x and Plan Apochromat IR 60x water-immersion; the acquisition system was controlled through NIS-Elements software (Version 6.10).

Immunofluorescence analysis was done on NIS-Elements Advanced Research software (Version 6.10.01). Colocalization analysis was performed using the Coloc2 plugin in Fiji.

### Statistical analysis

The statistical analysis performed in each experiment are detailed in the corresponding figure legend. Statistical analyses for individual organoid experiments and ELISAs were performed using GraphPad Prism 10 software. Statistics for differential gene expression analysis were performed using DESeq2 software, while statistics for differential protein expression analysis were performed using the DEP package from R. Finally, statistical analysis for GSEA was performed in the fgsea package in R.

### Data availability

All raw sequencing data are available through Annotare online resource: bulk RNAseq data on organoids treated with MSN at t=0, 20 min and 24 hours have been deposited in Annotare database under the accession code “E-MTAB-17035”. The bulk RNAseq data comparing healthy and tumor-associated SMT can be found on the same database under accession code “E-MTAB-17036”. Mass spectrometry data generated for conditioned media in this study can be found at the PRIDE partner[79] repository with the following dataset project accession identifier: PXD079264 (https://www.ebi.ac.uk/pride/archive/projects/PXD079264/privatereviewdataset, Token: 3xM1s7hQm7df). All other relevant data is provided within the Supplementary Information files.

## Acknowledgements

We thank the Cellular and Molecular Imaging Core Facility (CMIC), animal care facility at NTNU (CoMed), and PROMEC (proteomic analyses) core facilities at NTNU. CMIC is funded by the Faculty of Medicine and Health Sciences at NTNU, Central Norway Regional Health Authority and NALMIN II (NFR 322607). CMIC HPC computational infrastructure depends on Sigma2 project NN12203K (NAIBA - Norwegian AI for Bioimage Analysis). PROMEC is part of NAPI (RCN INFRASTRUKTUR-program 295910) and supported by Sigma2 Projects NN9036K and NS9036K for data storage and computational analysis. H.T.L is supported with grants from South-Eastern Norway Regional Health Authority (2024040), The Research Council of Norway (359632), the Norwegian Cancer Society and the Norwegian Breast Cancer Society (318232). Funding for this work was provided by the Research Council of Norway (‘Young Research Talent’ 326209 to M.M.-A) and the Norwegian Cancer Society (245170 to M.M.-A).

## Author contributions

M.M.-A. and R.Y. conceptualize the study; R.Y., L.S.L., M.D., A. B. S., A.M., and U.N. performed experiments and/or analyzed data; R.Y. and M.D. made figures, A.D.-S. developed image analysis tools, H.T.L. helped with bioinformatic analyses, M.B. performed LC/MS (lipidomics), and L.H. and A.S. provided proteomics expertise. M.M.-A. and M.J.O. supervised, R.Y. and M.M.-A. wrote the first draft, and all authors provided input to the final draft.

## Supplementary Figures

**Figure S1.**
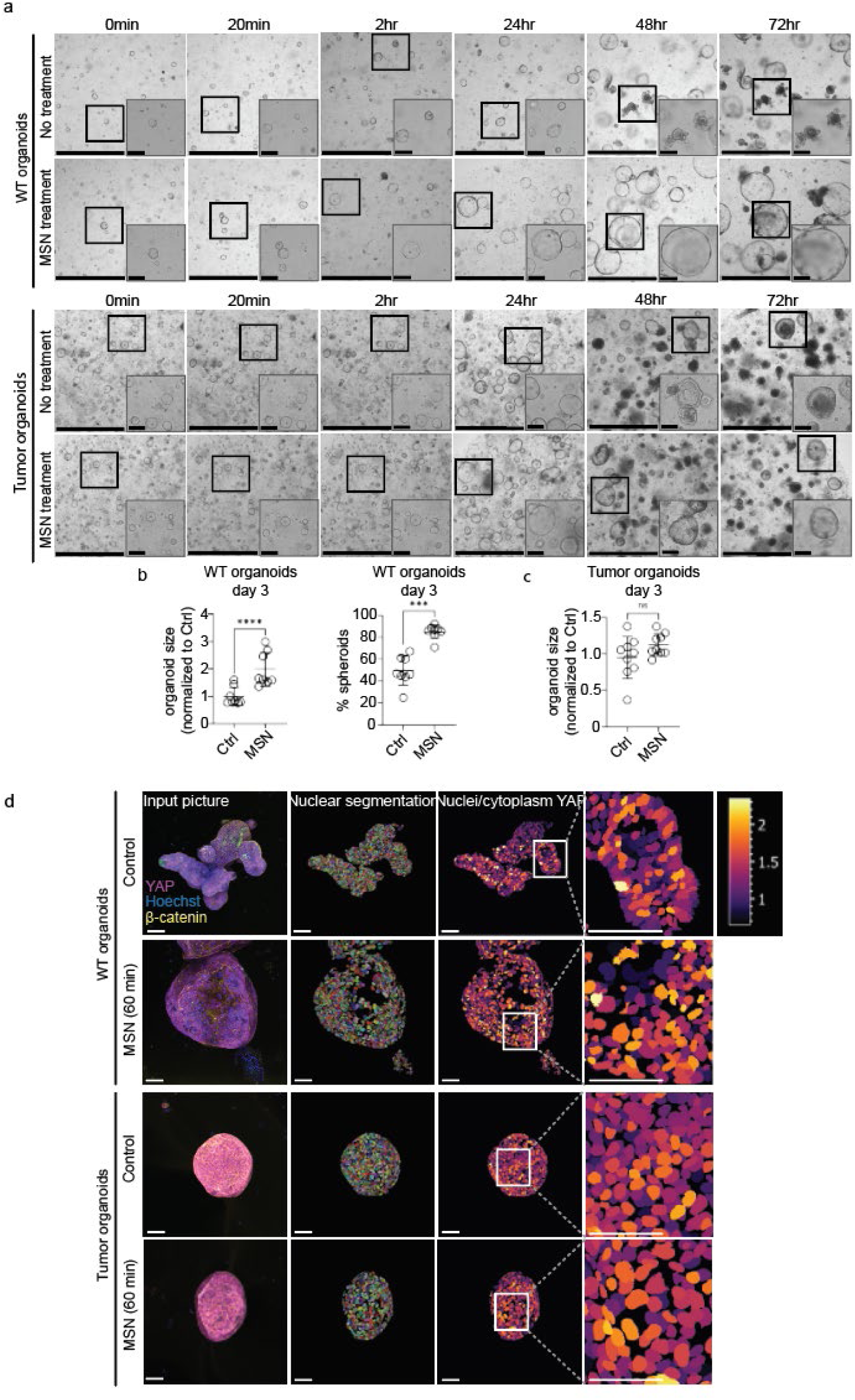
Timeline of organoids’ morphological changes in response to MSN and example of YAP signaling segmentation. **A.** Representative bright-field images showing organoids growth in response to Control vs MSN media at different timepoints (pretreatment t=0 until 72hours) in WT and Apc^Min/+^ tumor organoids. Scale bars = 1250µm and 200µm in the zoomed insets. **B.** Percentage of spheroids in organoid culture when grown in control or MSN, p=0.0001. **C.** Relative organoid size of Apc^Min/+^-derived tumor organoids at day 2 and day 3 post-seeding and treatment, p=0.008 and p=0.0710, respectively. **D**. Representative images of WT and tumor organoids cultured in control medium or treated with MSN for 60 min, illustrating YAP nuclear translocation analysis. Shown are YAP-stained organoids (as in Figure 1E), nuclear segmentation masks, and corresponding nuclear-to-cytoplasmic (N/C) YAP ratio colormaps. The colormap scale depicts relative N/C YAP ratio values. Scale bar = 50 µm for all images.

**Figure S2.**
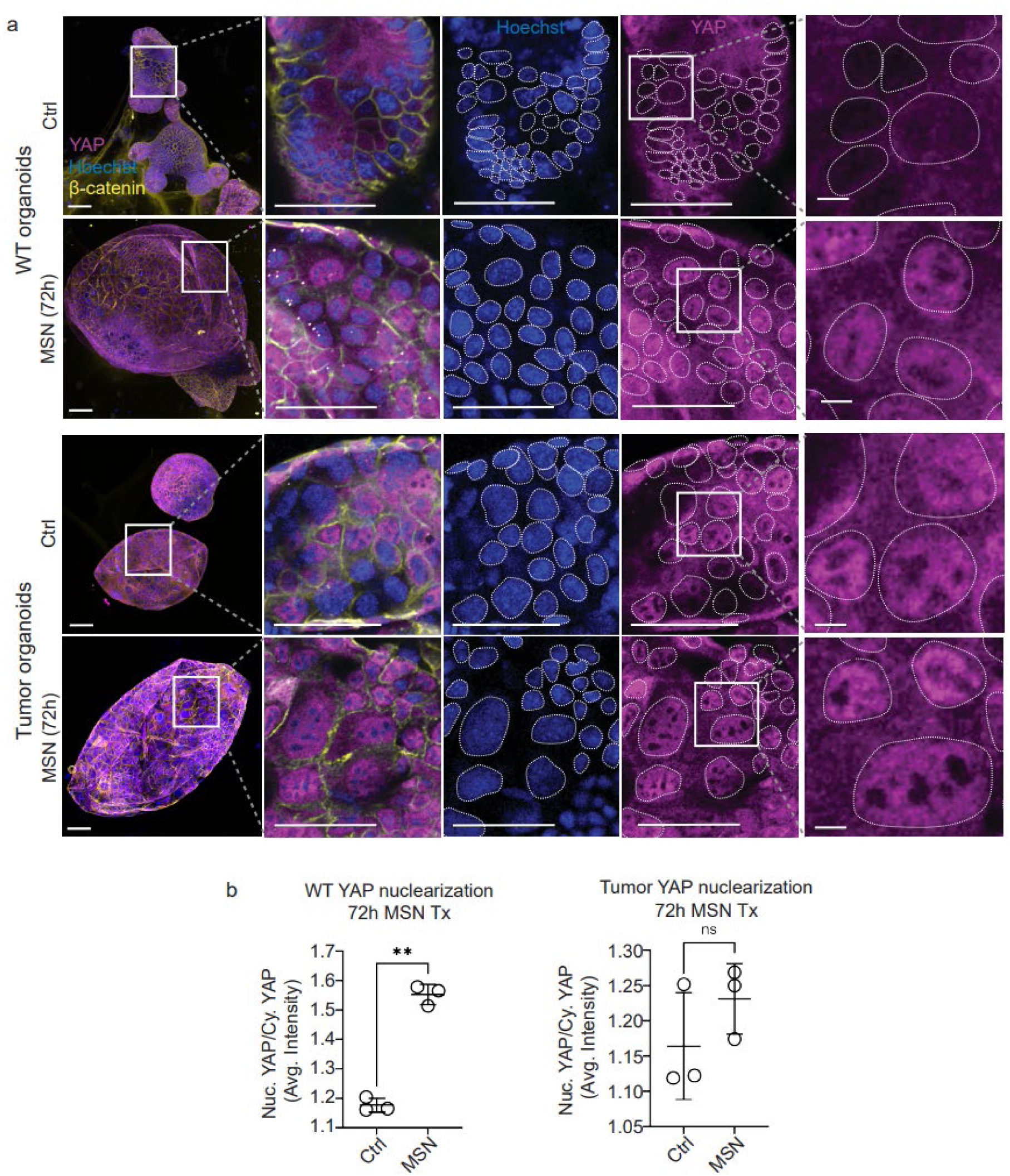
YAP subcellular localization in organoids after treatment with muscle supernatant for 72h and example of automated image analyses masks. **A.** Representative confocal immunofluorescence staining of YAP subcellular localization in WT organoids and tumor organoids after 72h treatment with MSN or control media. Scale bars represent 50µm in overview images and individual channel images and represent 5µm in the isolated zoomed YAP channels. **B.** Graphs represent relative subcellular YAP localization (proportion of nuclear YAP to cytoplasmic YAP) in WT and tumor organoids treated with MSN for 72h immediately upon seeding, as quantified by automated image analysis (n=3 biological replicates for both experiments, a total of 5-7 images were quantified per biological replicate). Paired t-test, p=0.0018 and p=0.4540 for WT and tumor organoids, respectively. Error bars depict mean values +/-standard deviations.

**Figure S3.**
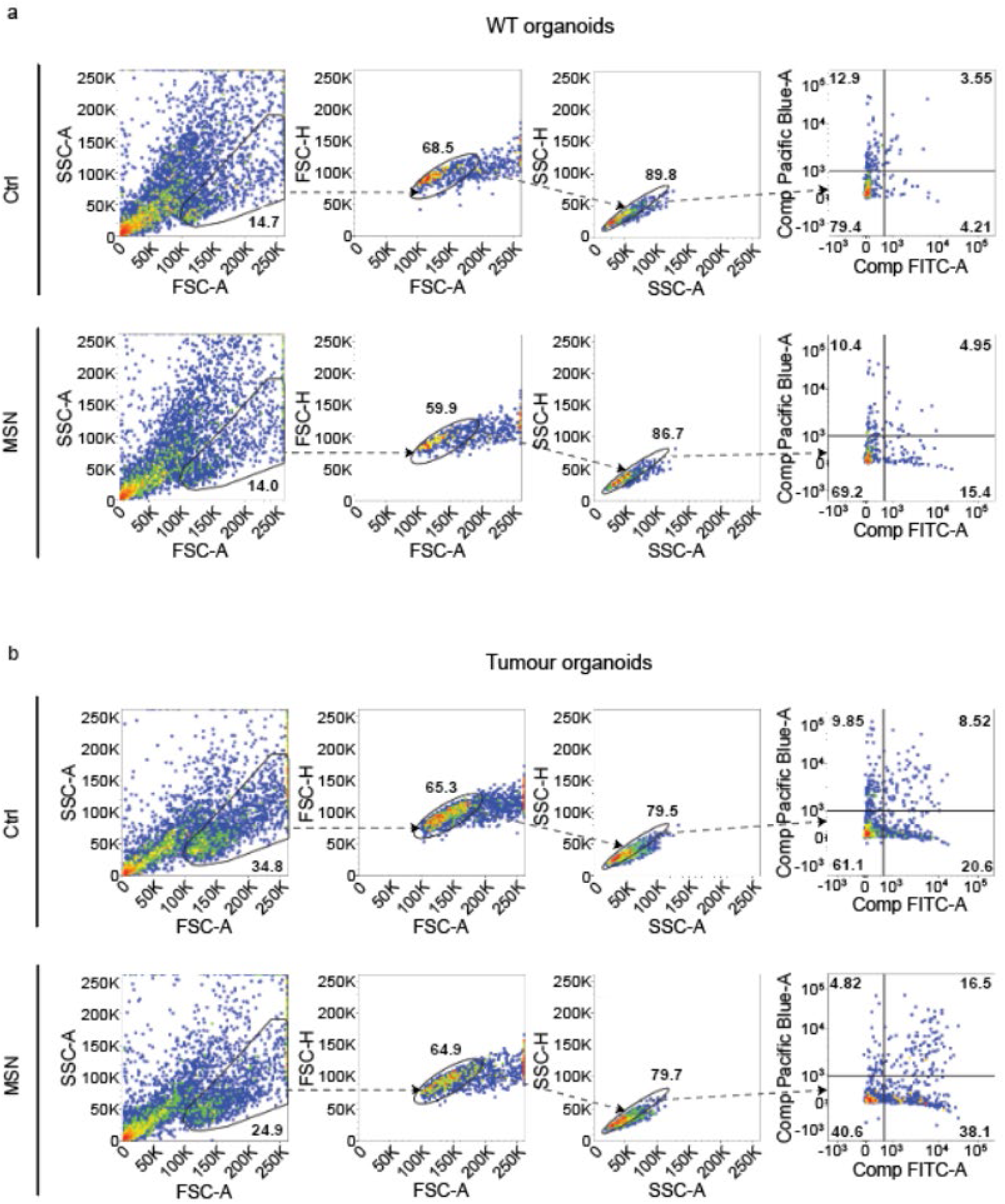
Flow cytometry gating strategy and example analysis for Sca1+ cells in control and MSN-exposed WT and tumor organoids. Charts show flow cytometry gating for intestinal organoids, where Pacific Blue represents dead cells stained with DAPI and FITC represents Sca-1+ expression. **A.** Example flow cytometry gating for WT organoids **B.** Example flow cytometry gating for tumor organoids.

**Figure S4.**
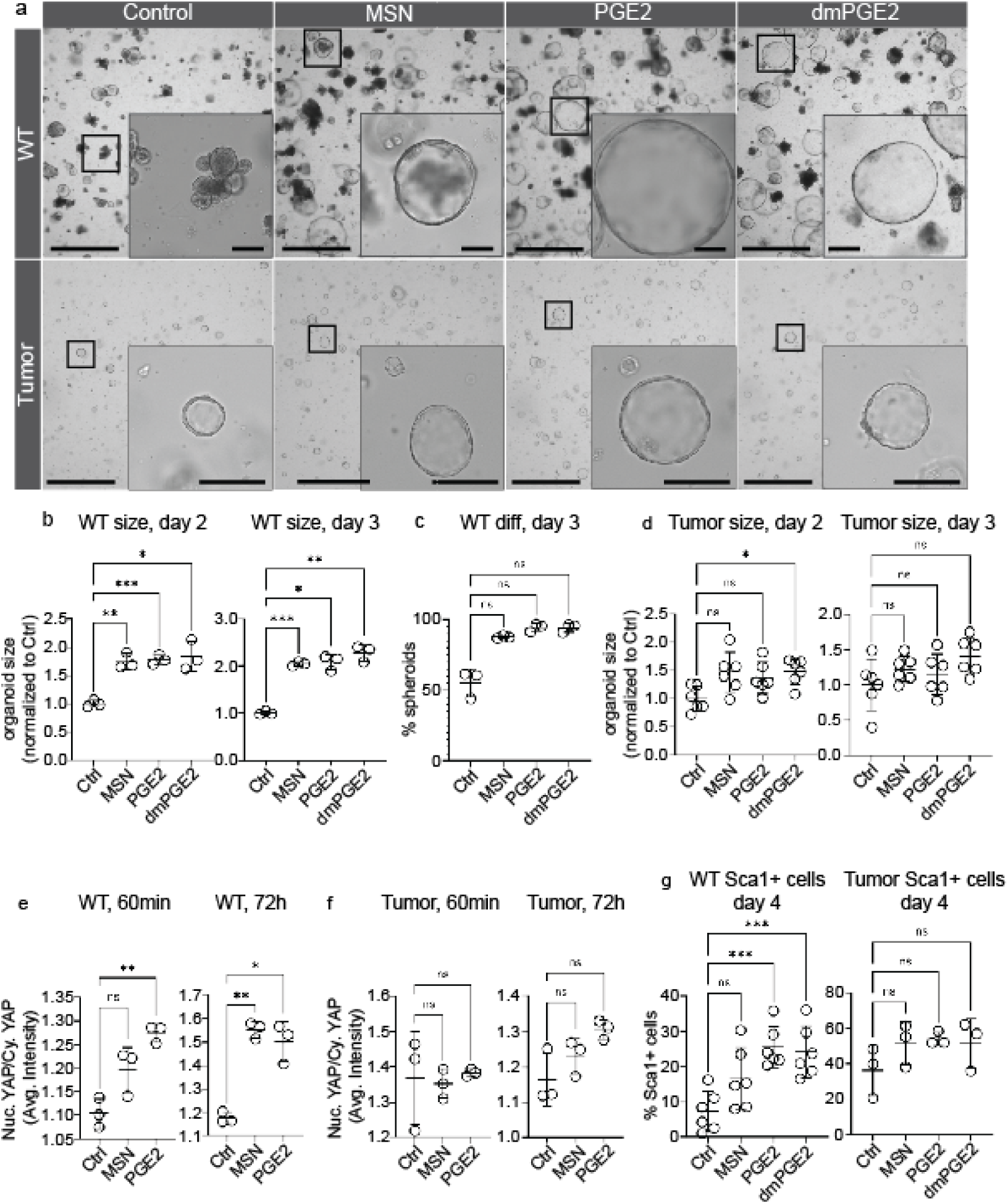
Organoid morphological response to PGE2 mimics MSN-induced phenotype in organoids. **A.** Representative bright-field images of WT and tumor organoids treated for 3 days with media supplemented with Control, MSN, PGE2, or dmPGE2. Scale bars represent 1250µm and 200µm in zoomed insets. **B-G**. Graphs show experiments on organoid cultures where points represent mean values from a separate organoid line. Repeated measures one-way ANOVA statistic was applied. Asterisks show significant results of Dunnett’s multiple comparisons test. **B.** WT organoid size at day 2 and day 3 post seeding when treated with exogenous Control, MSN, PGE2, dmPGE2, or Control. For day 2: p=0.0117; padj=0.0089, padj=0.0008, and padj=0.0375 and day 3: p=0.0059; padj=0.0002, padj=0.0116, and padj=0.0088 for Control v MSN, Control v PGE2, and Control v dmPGE2, respectively. n=3. **C**. Percentage of spheroids in WT organoid culture after treatment with Control, MSN, PGE2, and dmPGE2: p=0.0272; padj=0.0540, padj=0.0565, and padj=0.0534 for Control v MSN, Control v PGE2 and Control v dmPGE2, respectively. n=3. **D.** Tumor organoid size at day 2 and day 3 post seeding when treated with Control, MSN, PGE2, or dmPGE2. For day 2: p=0.0178; padj=0.0831, padj=0.0873, and padj=0.0130 and for day 3: p=0.0412; padj=0.3063, padj=0.7438, padj=0.1103 for Control v MSN, Control v PGE2, and Control v dmPGE2, respectively. n=6. **E,F.** Graphs show relative subcellular YAP localization (proportion of nuclear YAP to cytoplasmic YAP) in organoids treated with Control, MSN, or PGE2 for 60min at day 2 post seeding or treated with PGE2 for 72h immediately upon seeding, compared with organoids receiving no treatment. n=3. **E.** YAP nuclearization in WT organoids. For 60min treatment at day 2: p=0.0027; padj=0.0558 and padj=0.0073 and for 72h treatment from day 0: p=0.0082; padj=0.0028 and padj=0.0244 for Control v MSN and Control v PGE2, respectively. **F.** YAP nuclearization in tumor organoids. For 60min treatment at day 2: p=0.8; padj=0.9655 and padj=0.9772 and for 72h treatment from day 0: p=0.1882; padj=0.6305 and padj=0.1257 for WT v MSN and WT v PGE2, respectively. **G**. Sca1+ cell population for WT organoids; p=0.0026; padj=0.0588, padj=0.0005, and padj=0.0003 for Control v MSN, Control v PGE2, and Control v dmPGE2, respectively. n=6. **E. F.** Sca1+ cell population for tumor organoids; p=0.3141; padj=0.1372, padj=0.2210, and padj=0.5442 for Control v MSN, Control v PGE2, and Control v dmPGE2, respectively. n=3. All numerical graphs depict mean values +/-standard deviation.

**Figure S5.**
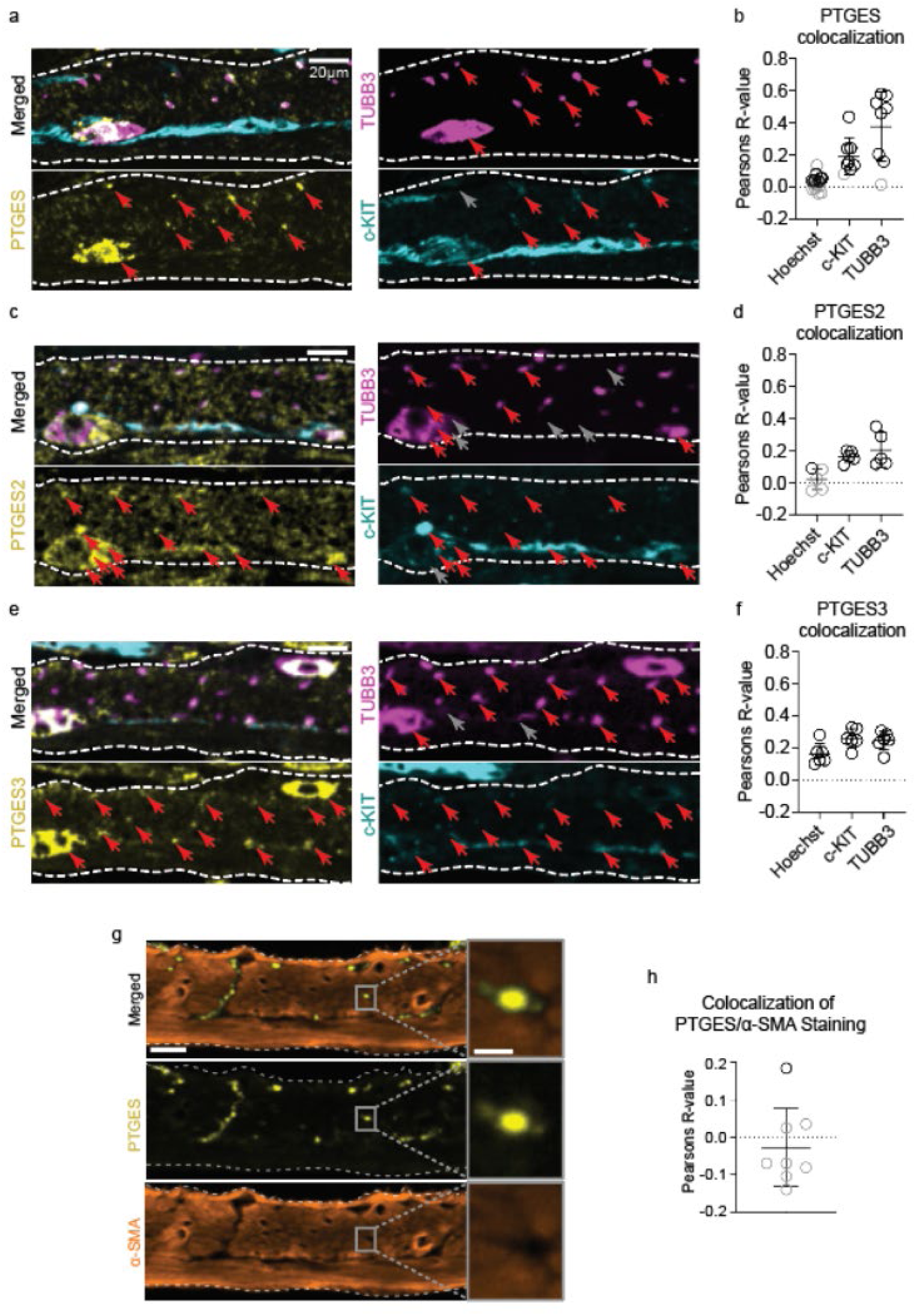
PGE2-converting enzymes PTGES, PTGES2 and PTGES3 expression patterns in the SMT. A, C, **E.** Immunostaining images show representative confocal images of SMT. PGE2-converting enzyme is shown in yellow (PTGES, PTGES2 and PTGES3, respectively), TUBB3 is shown in magenta, and c-KIT is shown in cyan. The area of the smooth muscle is delineated by white dotted lines. Red arrows mark areas of heightened expression in PGE2-converting enzyme and areas in corresponding panels with colocalized signal, whereas grey arrows mark areas in corresponding panels that are negative for colocalized signal. Scale bars shown in the top left panels represent 200µm and applies to all panels. **B. D. F**. Plots show the colocalization of PGE2-converting enzyme with Hoechst, c-KIT, and TUBB3 calculated using Coloc2 in Fiji. Each dot represents mean results of immunohistochemistry colocalization from a separate mouse, using data from 1-5 images per mouse. Black dots represent significant colocalization, whereas grey dots represent insignificant colocalization results. Error bars depict mean values +/-standard deviation. **B.** PTGES colocalization; n=16 for PTGES/Hoechst colocalization and n=8 for PTGES/c-KIT and PTGES/TUBB3 colocalization. **D**. PTGES2 colocalization; n=5 for all analyses. **F**. PTGES3 colocalization; n=6 for all analyses. **G**. Representative confocal images of immunohistochemistry staining of mouse intestinal SMT, showing localization of PTGES (yellow) in relation to α-SMA (orange). The area of the SMT is delineated by white dotted lines. Scale bars represent 20µm in wide-view images and 4µm in zoomed boxes. (n= 8 mice) **H.** Quantification of colocalization between PTGES and α-SMA using Coloc2 in Fiji. Each dot represents mean results from one separate mouse (n=8), using data from 1-5 images per mouse. Black dots represent significant colocalization, whereas grey dots represent insignificant colocalization. All numerical graphs depict mean values +/-standard deviation.

**Figure S6.**
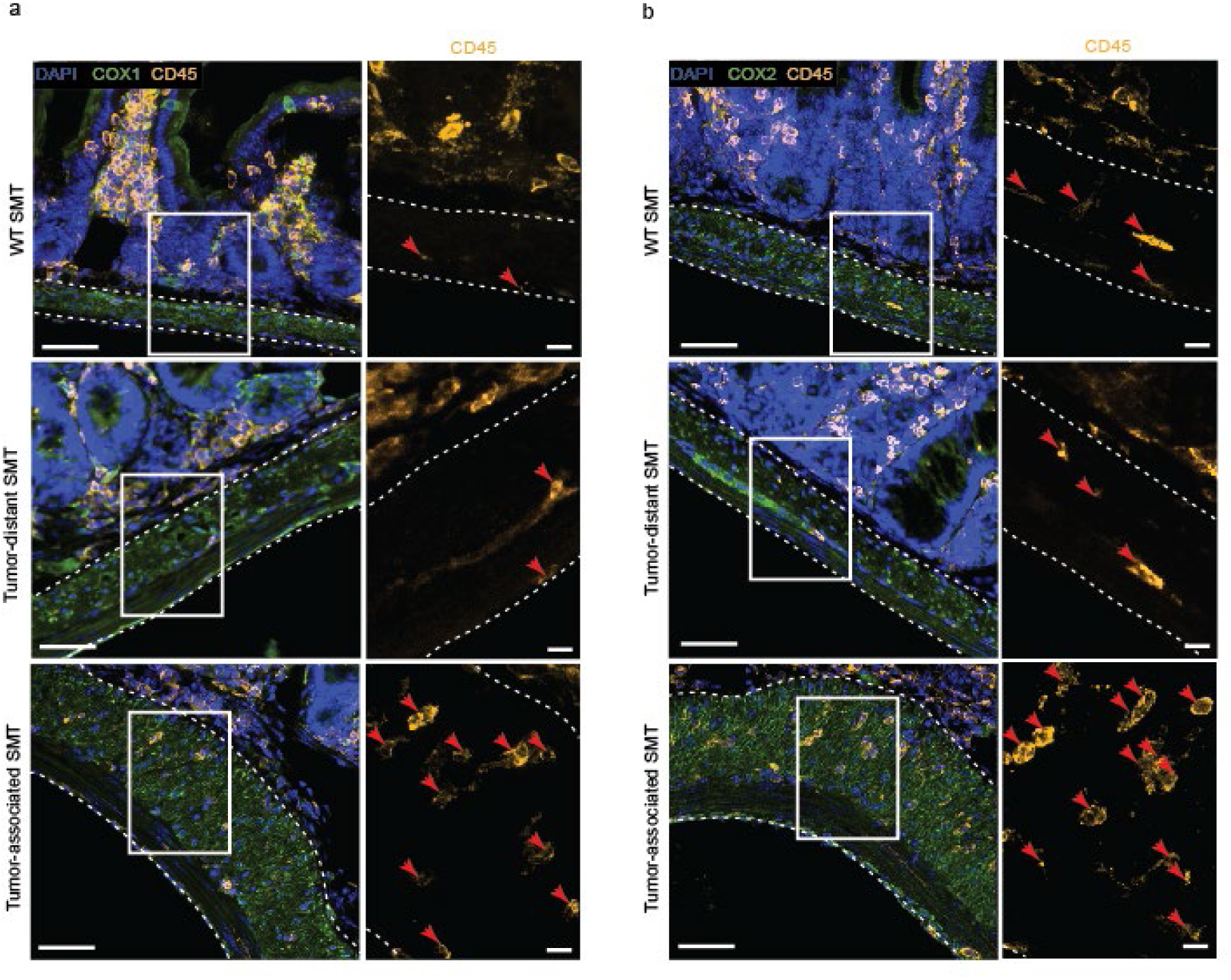
Immune infiltrate in the tumor-associated muscle expresses Ptgs1 and 2 (COX1 and 2). **A.** Representative confocal images of immunohistochemistry staining of mouse intestine showing COX1 (green) and pan-leukocyte CD45 marker (yellow) staining in WT, tumor-distant and tumor-associated SMT. The SMT is delineated by white dotted lines. Red arrows mark CD45+ cells in the SMT. Scale bars represent 50µm and 10 µm in zoomed insets. Note the increased CD45+ infiltrate, usually COX1 negative. **B.** Representative confocal images of immunohistochemistry staining of mouse intestine showing COX2 (green) and CD45 (yellow) staining in WT, tumor-distant and vs tumor-associated SMT, as in **A**.

**Figure S7.**
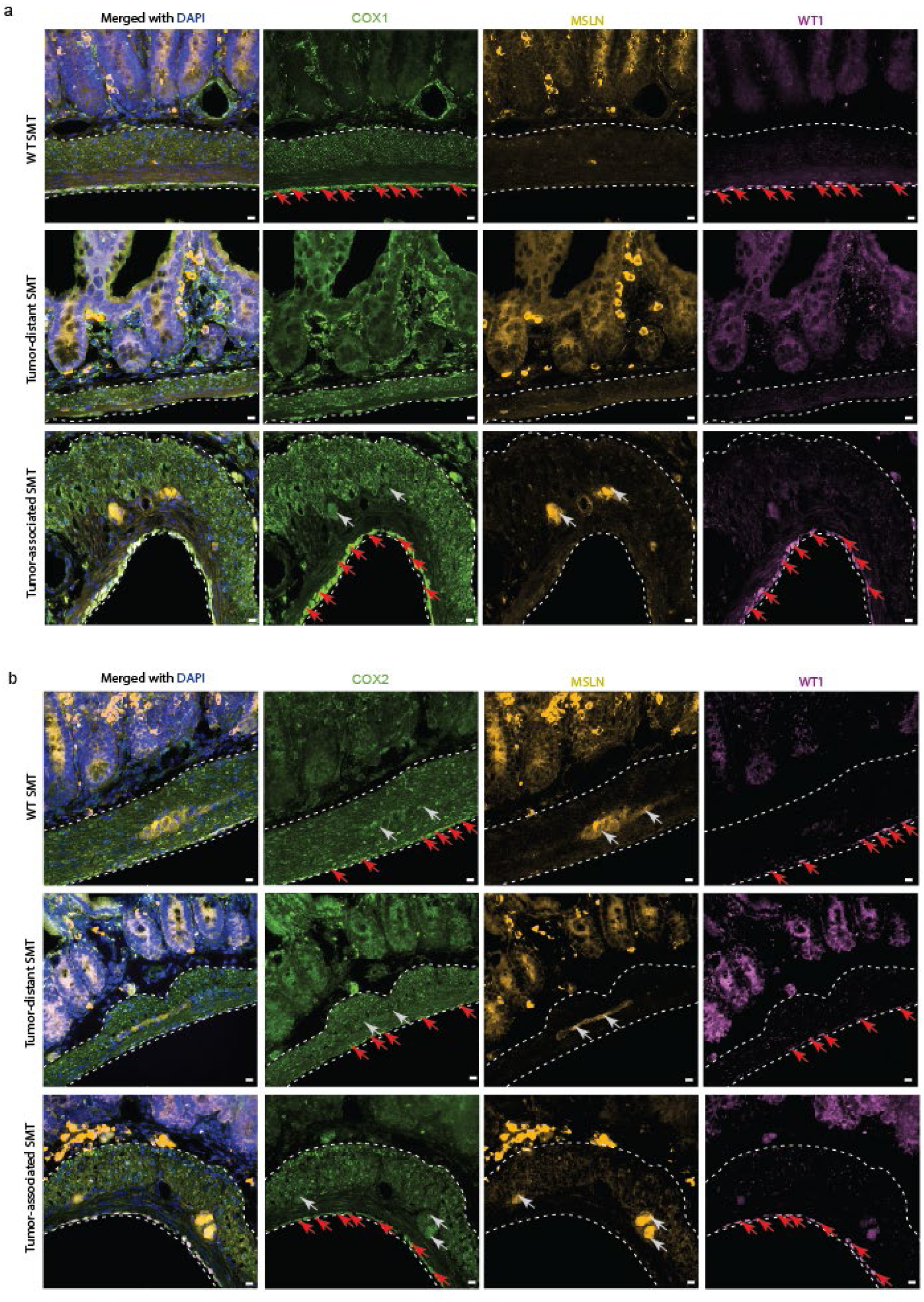
COX1 and 2 are expressed by mesenchymal cells in the SMT and serosa. A, **B.** Representative confocal images of WT and Apc^Min/+^ mouse intestine showing COX (**A**) or COX (**B**) staining (green) in WT, tumor-distant and tumor-associated SMT (from Apc^Min/+^ mouse) in relation to mesenchymal cell markers MSLN (yellow) and WT1 (pink). The SMT is delineated by white dotted lines. Grey arrows mark areas of heightened expression of MSLN and corresponding areas in COX1/2 panels, whereas red arrows highlight areas of heightened expression of WT1 and corresponding areas in COX1/2 panels. Scale bars represent 10 µm.

**Figure S8.**
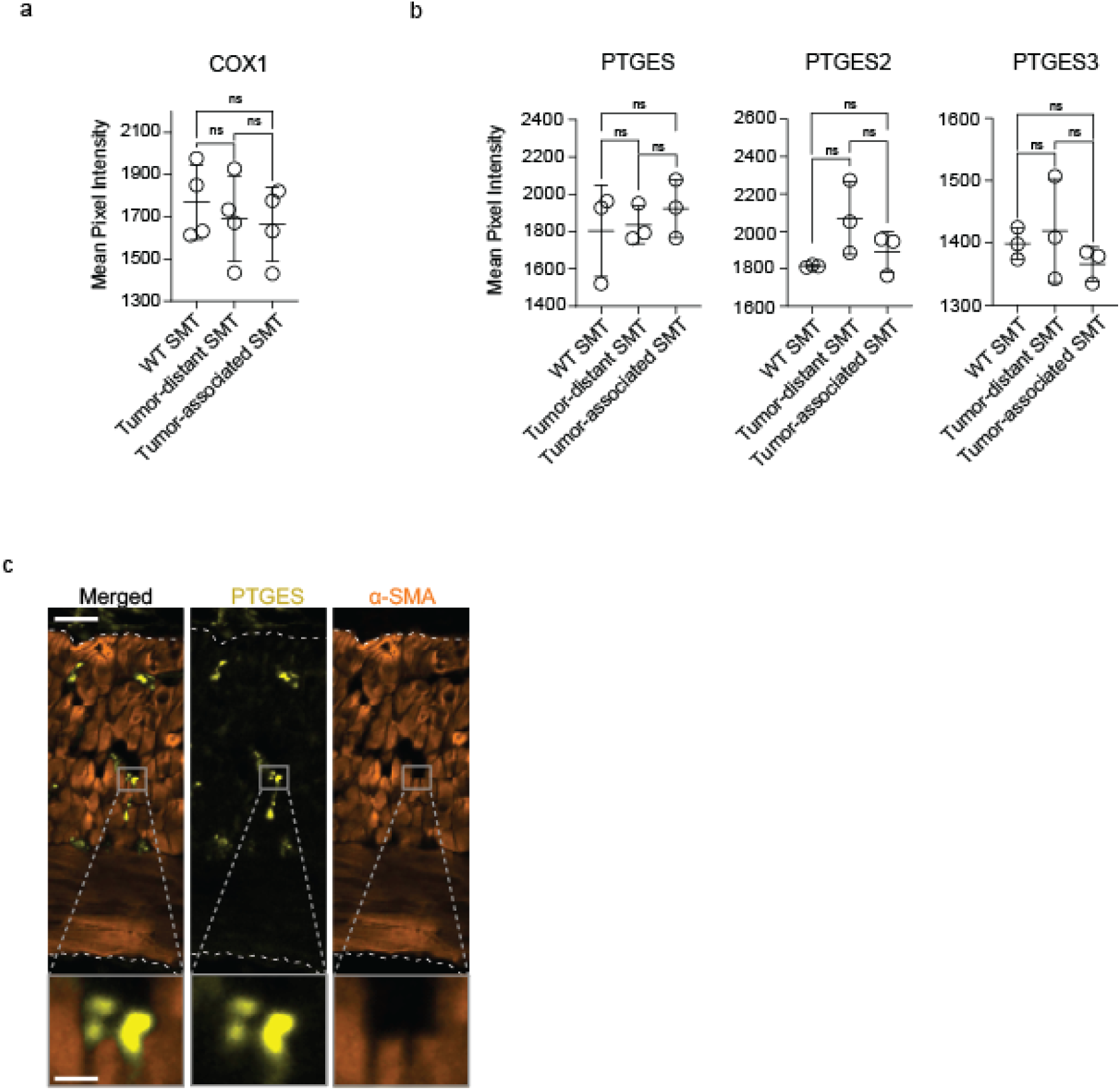
Further characterization of PGE2 converting enzymes expression patterns in SMT. **A.** Quantification of average COX1 intensity (per pixel) in WT, tumor-distant and tumor-associated SMT. Each dot represents the mean results of immunohistochemistry intensity measurement from a mouse, using data from 3-5 images per mouse (n = 4). Average COX1 intensity; WT SMT vs tumor-distant SMT; p=0.5912; WT SMT vs tumor-associated SMT; p=0.4411; and tumor-distant SMT vs tumor-associated SMT; p=0.4554. **B.** Quantification of average PGE2 synthase intensity (per pixel) in WT, tumor-distant and tumor-associated SMT. Each dot represents the mean results of immunohistochemistry intensity measurement from a mouse, using data from 3-5 images per mouse (n = 3). Average PTGES intensity; WT SMT vs tumor-distant SMT; p=0.8446; WT SMT vs tumor-associated SMT; p=0.5156; and tumor-distant SMT vs tumor-associated SMT; p=0.2757. Average PTGES2 intensity; WT SMT vs tumor-distant SMT; p=0.0854; WT SMT vs tumor-associated SMT; p=0.3011; and tumor-distant SMT vs tumor-associated SMT; p=0.1158. Average PTGES3 intensity; WT SMT vs tumor-distant SMT; p=0.7030; WT SMT vs tumor-associated SMT; p=0.2058; and tumor-distant SMT vs tumor-associated SMT; p=0.2671. All numerical graphs in **A.** and **B.** depict mean values +/-standard deviation. Statistical analyses were done with unpaired t-tests to compare WT and Apc^Min/+^ SMT and with paired t-tests to compare tumor-distant and tumor-associated SMT. **C.** Representative confocal images of immunohistochemistry staining of mouse intestinal tumor-associated SMT under epithelial tumor, showing localization of PTGES (yellow) in relation to α-SMA (orange). SMT area is delineated by white dotted lines. Scale bars represent 20µm in wide-view images and 4µm in zoomed boxes (n=3 ApcMin mice analyzed).

---

**Table S1.**
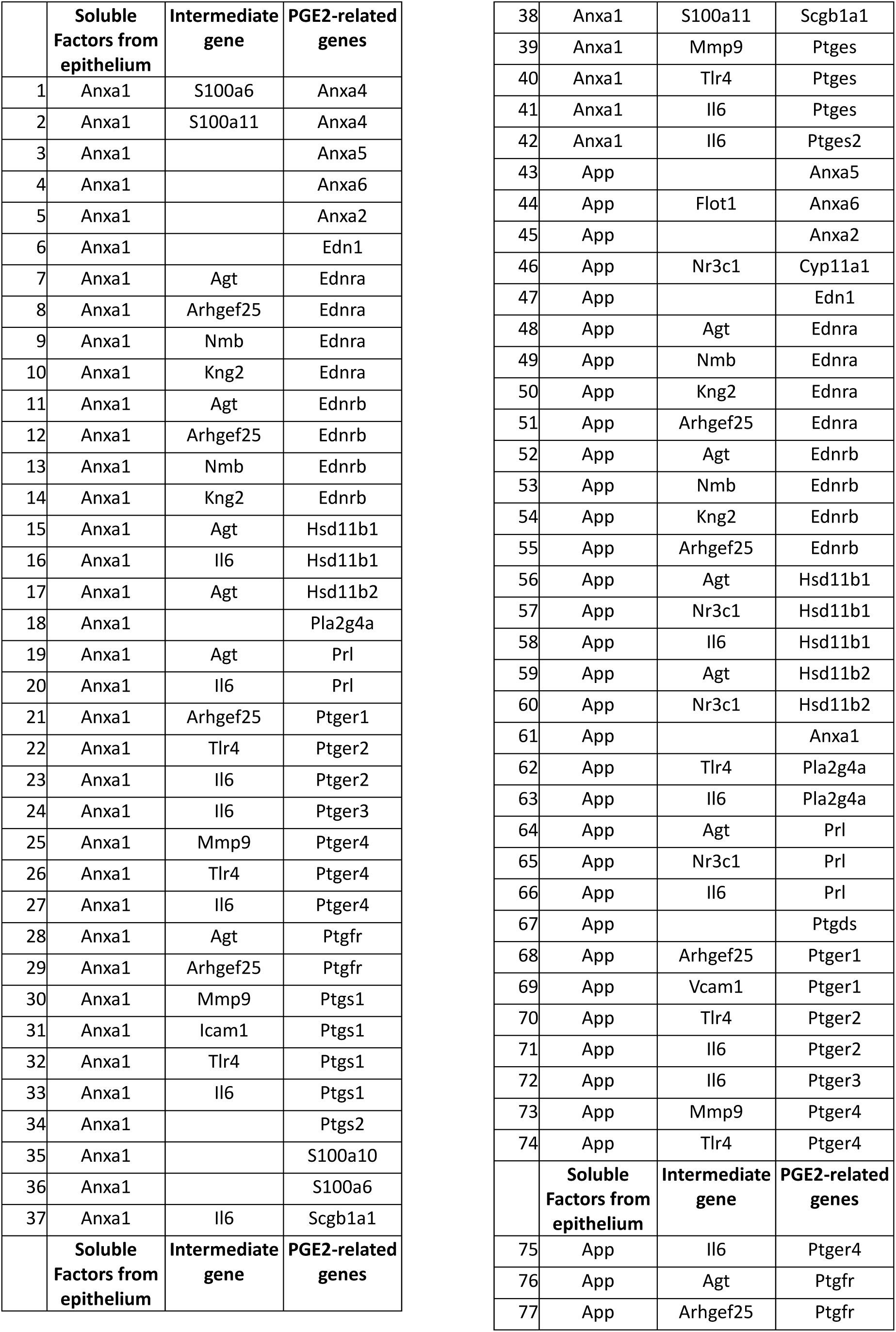
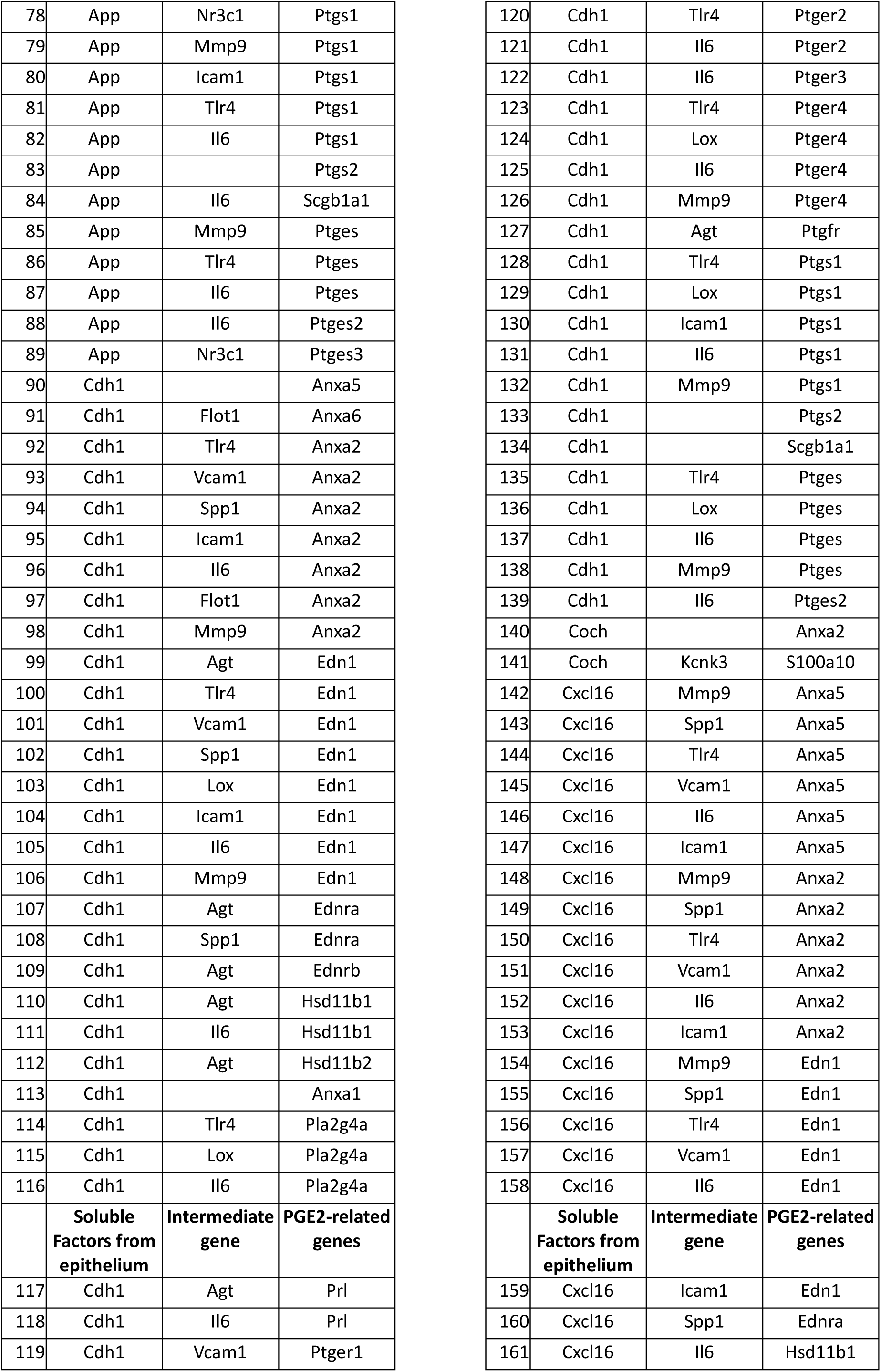
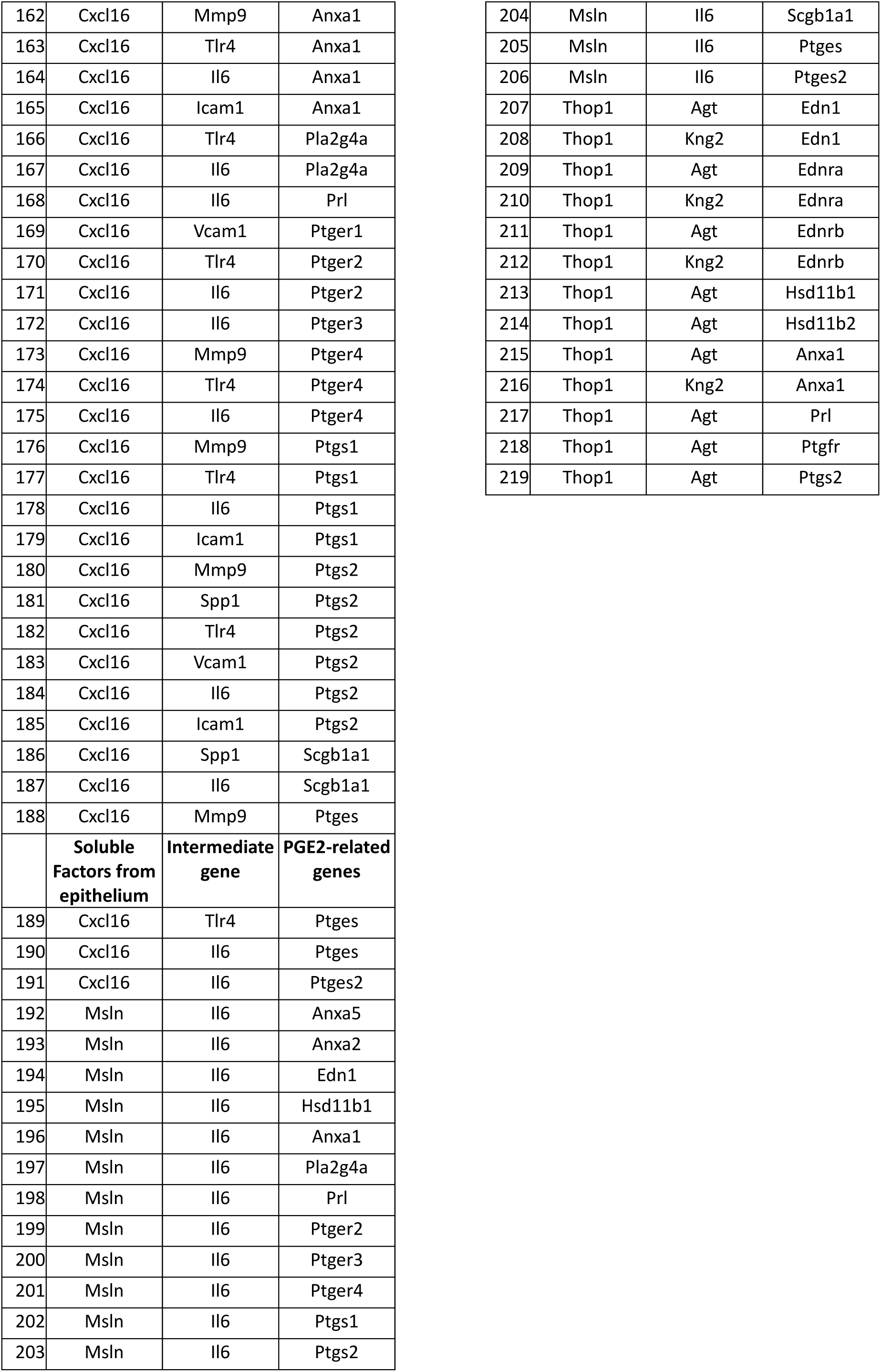
List of proposed interactions between soluble muscle factors and prostaglandin pathway genes or proteins, either directly associated or connected through ‘intermediate genes’, which were identified to be upregulated or downregulated in tumor-associated muscle.

## Notes

### Competing Interest Statement

The authors have declared no competing interest.

